# Repurposing screen identifies Amlodipine as an inducer of PD-L1 degradation and antitumor immunity

**DOI:** 10.1101/2020.05.26.117770

**Authors:** Chushu Li, Han Yao, Huanbin Wang, Jing-Yuan Fang, Jie Xu

## Abstract

Cancer cell expression of PD-L1 leads to T cells exhaustion by transducing co-inhibitory signal, and further understanding the regulation of PD-L1 in cancer cells may provide additional therapeutic strategies. Here by drug repurposing screen we identified amlodipine as a potent inhibitor of PD-L1 expression in cancer cells. Further survey of calcium-associated pathways revealed calpains-dependent stabilization of the PD-L1 protein. Intracellular calcium delivered an operational signal to calpain-dependent Beclin-1 cleavage, blocking autophagic degradation of PD-L1 accumulated on recycling endosome (RE). Blocking calcium flux by amlodipine depleted PD-L1 expression and increased CD8+ T cell infiltration in tumor tissues but not in myocardium, causing dose-dependent tumor suppression *in vivo*. Rescuing PD-L1 expression eliminated the effects of amlodipine, suggesting the PD-L1 dependent effect of amlodipine. These results reveal a calcium-dependent mechanism controlling PD-L1 degradation, and highlight calcium flux blockade as a potential strategy for combinatorial immunotherapy.

## Introduction

Programmed death-ligand 1 (PD-L1, also termed B7-H1 or CD274) is an immunosuppressive ligand which binds programmed cell death 1 (PD-1) on T cells to transduce inhibitory signal ^1^. The expression of PD-L1 is required for the maintainance of physiological immunotolerance, but cancer cells may ectopically express PD-L1 to evade immune surveillance. Antibody inhibitors of PD-1/PD-L1 have displayed considerable therapeutic effects to an increasing number of solid tumors ^2^, but still facing challenges such as low response rate, acquired resistance after long-term treatment, and immune-related adverse effects (irAEs) ^3 4^. A considerable fraction of PD-L1 protein is constantly internalized, recycled and redistributed between the cell membranes and recycling endosomes^5^, which may facilitate the escape from being neutralized by antibody drugs. To increase the efficacy of PD-L1 blockade, it is a plausible strategy to target the whole-cell PD-L1 by selective degradation.

In previous studies, we and other groups have revealed pathways involved in PD-L1 degradation, including the endocytosis and lysosomal degradation mediated by HIP1R^6^, the endoplasmic reticulum associated degradation (ERAD) by de-glycosylation or AMPK blockade ^7^, as well as autophagic degradation ^8 9^. As a type-I transmembrane protein, PD-L1 is expected to be regulated by multiple factors in the protein production, modification and destruction processes. However from a therapeutic perspective, a more important question is to achieve tumor-specific deplection of PD-L1 by approaches with limited adverse effects. The discovery of new indications for existing medications, known as drug repurposing, provides an attractive alternative to de novo drug development.

In a repurposing screen, we found calcium channel blocker Amlodipine potently induced the degradation of PD-L1 but not other tested proteins such as major histocompatibility complex (MHC), suggesting the involvement of selective autophagic mechanisms. Currently, our understanding on the selective autophagy is still limited. It is proposed that autophagosome membranes may derive from endoplasmic reticulum (ER) ^10-12^, mitochondria ^13 14^, the Golgi apparatus ^15 16^, the recycling endosomes ^17-19^ and plasma membrane ^20^. In addition to non-selective sequestration of cellular components usually induced by nutritional deprivation, increasing evidence has suggested that autophagy can also be responsible for the removal of specific cellular materials, such as protein aggregates (aggrephagy), aberrant mitochondria (mitophagy), pathogens (xenophagy) and other superfluous organelles ^21-25^. The mechanisms of selective autophagy are intricate and not yet fully understood. Interestingly, the recycling endosome (RE) was recently found to provide the origin membrane and platform for the autophagosomes ^26^. However, it is unknown whether selective autophagy of RE may occur, and if so, how it could be regulated in cancer cells.

In the present study, we show that amlodipine triggers selective autophagy of RE, thereby causing the degradation of PD-L1 that is highly enriched on RE. We also show that amlodipine functions by blocking calpain-dependent cleavage of Becklin-1, which leads to RE autophagy. Importantly, the effects of amlodipine on promoting PD-L1 degradation displayed potent anti-tumor effect in a MC38 model in vivo, highlighting an alternative apporoach to selectively degrade PD-L1 for promoting anti-tumor immunity.

## RESULTS

### Identification of Amlodipine as a suppressor of PD-L1 expression

Our work started with drug repurposing by screening a clinical compound library (Selleck-Pfizer-Licensed-Library), attempting to find potential inhibitors of PD-L1 and improve the efficacy of PD-L1-associated immunotherapy. The brief description of the experiment process is displayed in Fig.1a. After excluding the drugs that had been reported to be related to PD-L1, such as the inhibitors of MEK^27-29^, FAK^30-34^ and ALK^35-37^, we found that amlodipine, a dihydropyridine calcium channel blocker^38^, abundantly decreased the expression of PD-L1 in cancer cells (Supplementary Fig. 1a). When human colorectal cancer (CRC) RKO cells and breast cancer MDA-MB-231 cells were treated with amlodipine in dose and time dependent manners, the expression of PD-L1 gradually decreased in both cancer cells (Fig. 1b and 1c). Amlodipine also took effect in other human cancer cells highly expressing wild-type PD-L1, such as LoVo CRC and A375 melanoma cancer cells (Supplementary Fig. 1b). Consistently, immunofluorescence showed that incubation with amlodipine resulted in down-regulation of PD-L1 proteins compared with the control (Fig.1d). Since amlodipine is a calcium channel blocker (CCB) widely used to lower blood pressure, we asked whether it suppressed PD-L1 by calcium flux blocking. Amlodipine is composed of a mixture of (R)- and (S)-amlodipine enantiomers, and only the latter structural possesses L-type calcium channel blocking function^39^. To this end, we treated RKO and MDA-MB-231 cells with Levamlodipine, a L-type amlodipine, and found PD-L1 was obviously reduced in both cell lines (Fig.1e and Supplementary Fig.1c). To further confirm the inhibitory effect of cytosolic calcium on PD-L1, another calcium channel blocker nifedipine and the Ca2+ chelator BAPTA were respectively treated in tumor cells, both of which were able to suppress the expression of PD-L1 detected by western blot (Fig.1f-g and Supplementary Fig.1d-e). 2,5-Di-t-butyl-1,4-benzohydroquinone (BHQ) is a SERCA inhibitor and thapsigargin blocks the sarco-endoplasmic reticulum Ca^2+^ ATPase, both of which can elevate the concentration of calcium in cytoplasm. As expected, both drugs augmented the expression of PD-L1 (Fig.1h-i and Supplementary Fig.1f-g). Intracellular calcium tested by fluorescent calcium binding dye Fluo-8 showed amlodipine and BAPTA abundantly decreased the concentration of cytoplasmic calcium, while BHQ and thapsigargin increased the cytoplasmic calcium concentration compared with the control respectively (Fig.1j). To ascertain the specific effect of amlodipine on PD-L1, we also detected the expression of MHC Class I (MHC-I) simultaneously. Interestingly, amlodipine did not impair the expression of MHC-I (Fig.1b and 1c). These data indicate that amlodipine selectively decreases PD-L1 expression in various cancer cells by downregulating cytosolic calcium.

**Fig. 1.**
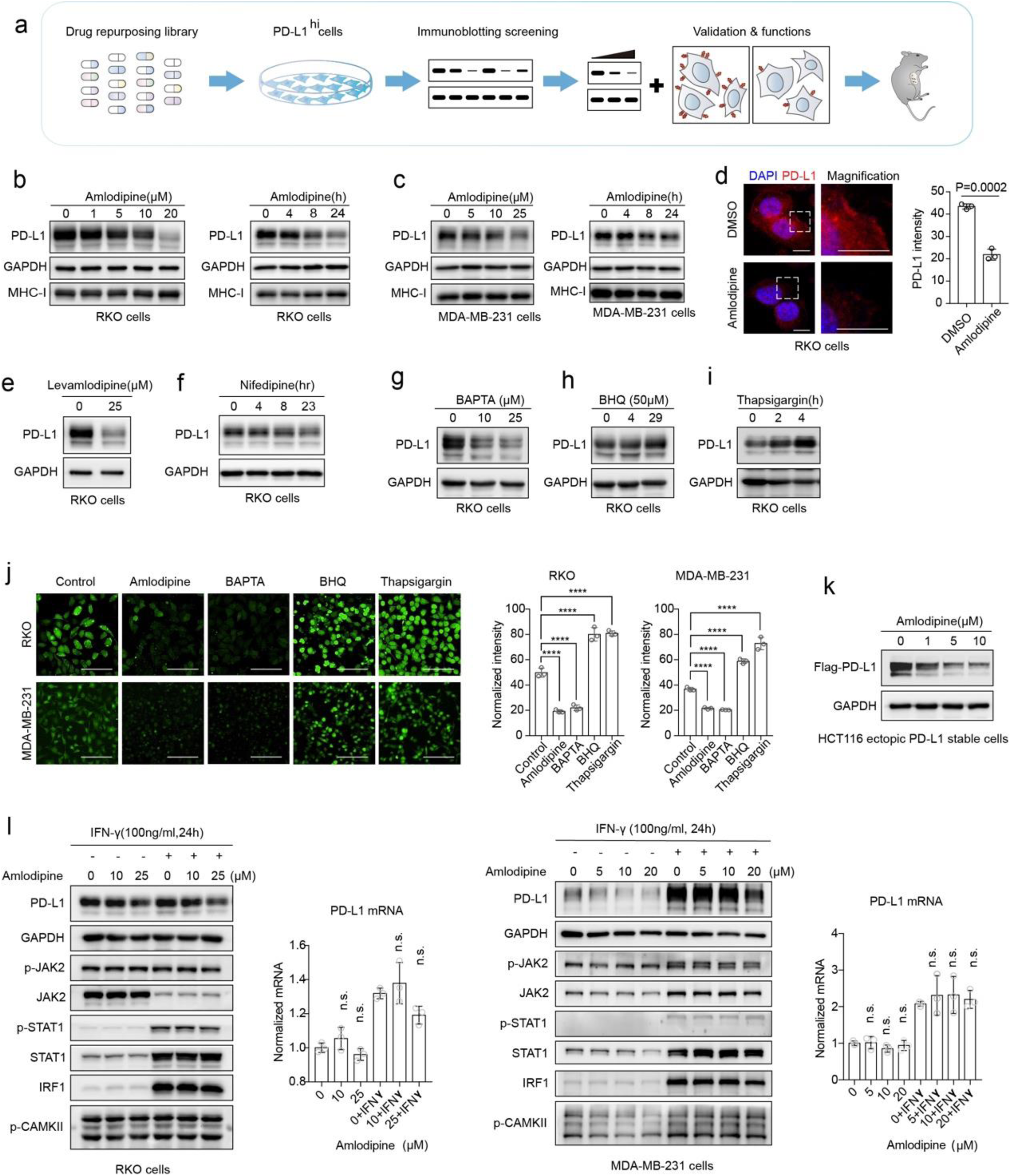
Amlodipine decreased PD-L1 expression post-translationally by blocking intracellular calcium flux. **a**, Brief description of our work process for the drug screening and validation. **b**,**c** RKO cells (**b**) and MDA-MB-231 cells (**c**)were treated with amlodipine at the indicated concentrations for 24h or incubated with 25μM amlodipine for 4–24 h, detected by immunoblotting with PD-L1, GAPDH and MHC-I antibodies. These experiments were repeated three times independently with similar results. **d**, RKO cells were treated with amlodipine (25μM, 24h). Left, immunofluorescence assay, PD-L1 was marked red and the cell nucleus was colored blue with 4′,6-diamidino-2-phenylindole (DAPI). Scale bars indicate 10 μM. Dashed frames indicate the representative fields to be presented in magnification. Right, quantification of PD-L1 intensity. Values are means ± s.d. from *n* = 3 independent experiments. Statistical differences were determined by two-sided Student’s *t*-test. **e**, RKO cells treated with leamlodipine at the indicated concentrations for 4h. **f**, RKO cells treated with nifedipine at the indicated concentrations for 24h. **g**, RKO cells treated with BAPTA at the indicated concentrations for 3h. **h**, RKO cells treated with BHQ at the indicated concentrations for 4h. **i**, RKO cells treated with 500nM thapsigargin for 0-4 h. **e**-**i** were repeated three times independently with similar results. **j**, RKO cells and MDA-MB-231 cells incubated respectively with 25μM amlodipine, 10μM BAPTA, 50μM BHQ and 500nM thapsigargin or dimethyl sulfoxide (DMSO) as a control. Fluorescent detection of the intracytoplasmic calcium with fluo-8 staining. Scale bars, 100μm. Right, values are means ± s.d. from *n* = 3 independent experiments. Statistical differences were determined by ANOVA post-hoc test. ****P < 0.0001. **k**, HCT116 exogenous PD-L1 stable cells treated with amlodipine at the indicated concentrations for 24h, detected by western blot with PD-L1 and GAPDH antibodies. This experiment was repeated three times independently with similar results. **l**, RKO cells and MDA-MB-231 cells treated with amlodipine at the indicated concentrations and 100ng/mL IFN-γ for 24h, subjected to immunoblotting and RT-qPCR. Values are means ± s.d. from *n* = 3 independent experiments. The *P* value was determined by ANOVA post-hoc test. n.s, no significance.

Next, we probed how amlodipine suppressed PD-L1. As previously reported, IFN-γ is a potent inducer which substantially promotes the transcription of PD-L1 through JAK/STAT/IRF1 axis ^40^. The mRNA and protein levels of PD-L1 were respectively detected in RKO and MDA-MB-231 cells with amlodipine treatment in the presence or absence of IFN-γ. Simultaneously, the activity of the JAK2/STAT1/IRF1 pathway was tested. Western bolt and RT-PCR revealed that amlodipine attenuated the up-regulation of PD-L1 stimulated by IFN-γ in protein level rather than mRNA level (Fig.1l). Besides, there was no significant difference in the JAK/STAT/IRF cascade in cancer cells treated with amlodipine. Additionally, no obvious change of p-CAMKII was detected after amlodipine treatment compared with the abundant decrease of PD-L1. Similar results were obtained with amlodipine treatment in other cancer cells (Supplementary Fig.1h-j) or with the treatement of other calcium channel blocker like nifedipine (Supplementary Fig.1k). Then, we used HCT116 CRC cells which had very low endogenous PD-L1 expression to establish a cell line stably overexpressing flag-tagged PD-L1, in which ectopic PD-L1 was also suppressed by the treatment with amlodipine (Fig.1k). Thus, we speculated the effect of amlodipine on PD-L1 was depend on the blocking of intracellular calcium in the protein level rather than the mRNA level through IFN-γ/JAK2/STAT1/IRF1 pathway.

### Amlodipine induced degradation of PD-L1 but not MHC-I

Although amlodipine had no effect on PD-L1 mRNA, it indeed accelerated the degradation of PD-L1 (Fig.2a). Studies have disclosed that calcium is a strong regulator of autophagy ^41 42^, and L-type CCBs have been identified to induce autophagy in a mTOR-independent manner^43^. Therefore, we hypothesized that as a canonical L-type CCB, amlodipine facilitated the autophagic degradation of PD-L1. With amlodipine incubation, decreased PD-L1 and increased LC3-II were detected not only in RKO and MDA-MB-231 cells (Fig.2b), but also in HCT116 cells transiently or stably transfected with ectopic flag-PD-L1 (Fig.2d and Supplementary Fig.2a), remarkably, with no corresponding fluctuation of MHC-I expression (Fig.2b and d). Analysis of immunofluorescence showed that with amlodipine treatment, LC3 was notably activated and recruited in the cytoplasm, consistently with decreased PD-L1 on the cell membrane and increased co-localization between PD-L1 and LC3-labelled autophagosome (Fig.2c).

**Fig. 2.**
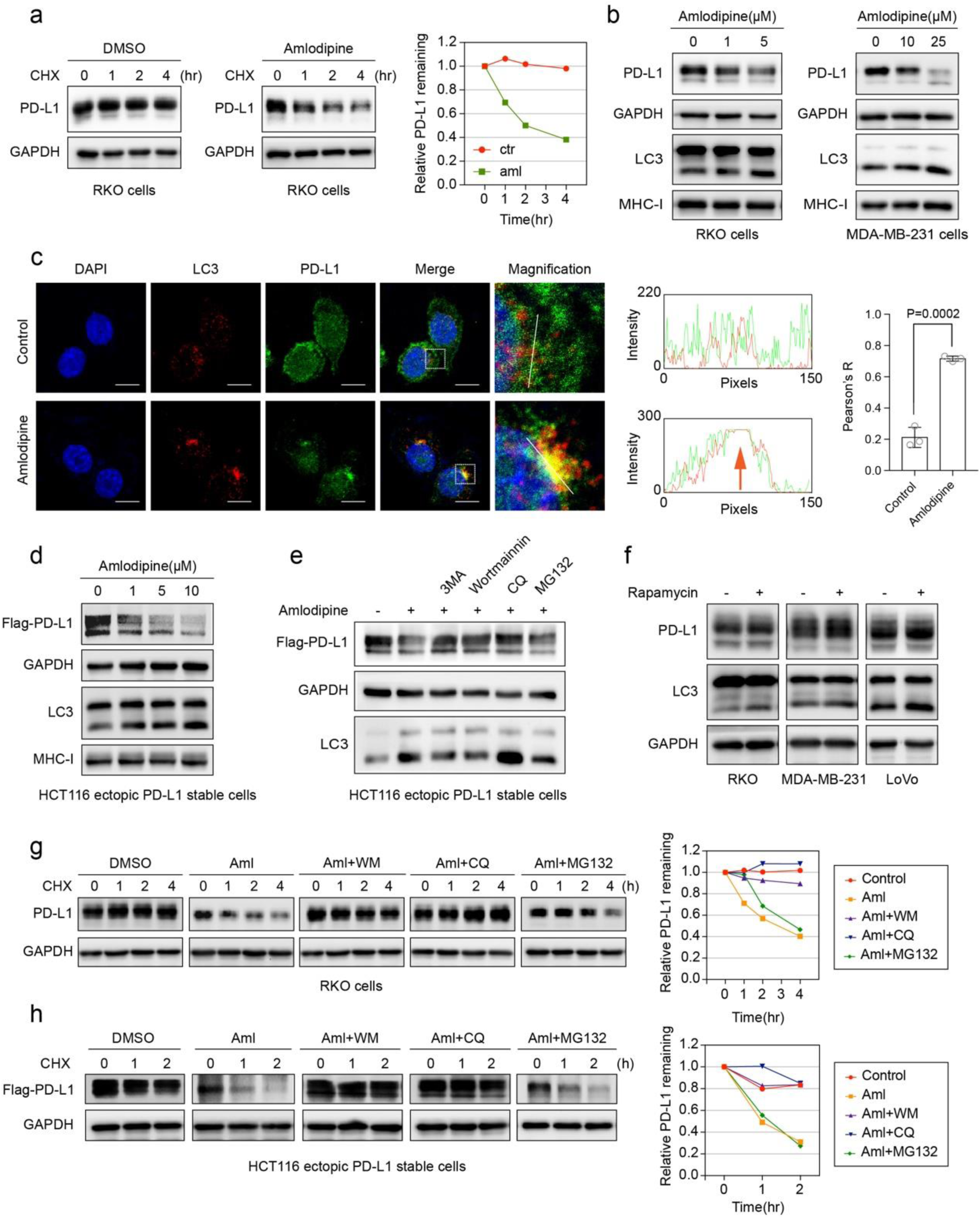
Amlodipine facilitated the autophagic degradation of PD-L1. **a**, RKO cells pretreated with DMSO or amlodipine(20μM) for 20h were co-incubated with cycloheximide (CHX) for indicated time points. Left, the levels of PD-L1 and GAPDH tested by western blot. Right, quantification of normalized remained PD-L1. This experiment was repeated three times independently with similar results. **b**,**d**, Immunoblots showing the effect of amlodipine on PD-L1, LC3 and MHC-I expression in the indicated cells. These experiments were repeated three times independently with similar results. **c**, Left, immunofluorescence showing the co-localization between PD-L1 and LC3 after amlodipine treatment. Scale bars, 10μm. Dashed frames indicate the representative areas to be presented in magnification. Middle, the intensity profiles of PD-L1 and LC3 along the white line, with the site of co-localization indicated by orange arrows. Right, values are means ± s.d. from *n* = 3 independent experiments. The P value was determined by two-sided Student’s *t*-test. **e**, Immunoblots showing the effect of amlodipine (20μM, 24h) on PD-L1 and LC3 expression in the absence or presence of autophagic inhibitors 3-Methyladenine (3-MA) and wortmannin(WM), lysosomal inhibitor chloroquine (CQ) or proteasomal inhibitor MG-132. **f**, The indicate cells treated with rapamycin(100nM) for 16h. **g**,**h**, Left, the degradation rate of PD-L1 detected at the indicated time points by CHX-chase assay in the indicated cells pretreated with DMSO or amlodipine in the presence of wortmannin(WM), chloroquine (CQ) and MG132 respectively. Right, quantification of the level of remained PD-L1. These experiments were respectively repeated three times independently with 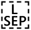 similar results.

Since the accumulation of LC3-II can be both detected owing to activated formation of autophagosomes or blocked fusion of autophagosome and lysosome, we used diverse small molecular inhibitors to further investigate the mechanism. The Vps34 inhibitors, 3-methyladenine (3-MA) and wortmannin, inhibit the formation of PI3P which therefore block the initiation of autophagy activity. Comparatively, chloroquine (CQ) blocks the distal process of autophagy by inhibiting the fusion of autopagosome and lysosome. The experiment revealed that the decrease of PD-L1 caused by amlodipine treatment could be rescued by co-incubation of 3-MA, wortmannin and CQ respectively, but not the proteasomal inhibitor MG132 (Fig.2e and Supplementary Fig.2b). By respectively blocking endogenous wild-type PD-L1 and ectopic flag-PD-L1 with amlodipine treatment, and co-incubating the tumor cells with different inhibitors of according degradation pathways, we used cycloheximide (CHX) to detect the degradation rate of PD-L1 proteins. Strikingly, wortmannin and CQ, rather than MG132 could attenuate the amlodipine-induced elimination of PD-L1 (Fig.2g and h). These data suggested that amlodipine facilitated the PD-L1 degradation by triggering autophagy flux.

Next, we questioned whether the autophagic degradation of PD-L1 could be triggered by other autophagy inducer. Interestingly, co-incubation with mTOR inhibitor rapamycin (Fig.2f) which facilitated bulk macroautophagy did not impair the expression of PD-L1 protein. Collectively, these findings indicated that amlodipine can accelerate some kind of selective autophagy of PD-L1.

### Amlodipine inhibits calpain-dependent stabilization of PD-L1

To further explore the correlation between decreased intracellular calcium flux and increased autophagic degradation of PD-L1 triggered by amlodipine, we started to look for an intermediate regulator between them. Calpains are a family of canonical calcium-associated cytosolic proteases, whose activity depends on the concentration of intracellular calcium^44^. Calcium-activated calpains function as proteolytic enzymes exerting protein cleavage. Previous studies have proved that calpains take part in CCB-induced autophagy ^42 43^. Western blot revealed that ectopic expression of flag-tagged calpain markedly suppressed the formation of LC3-II and increased the expression of both endogenous and exogenous PD-L1 (Fig.3a-b). By contrast, silencing calpain by specific siRNAs decreased the expression of PD-L1 and increased LC3-II without altering MHC-I expression (Fig.3c and Supplementary Fig.3a). Depletion of calpain also alleviated the enhancement of IFN-γ-induced PD-L1 (Supplementary Fig.3b). Consistently, incubation with calpeptin, a selective inhibitor of calpains, facilitated the down-regulation of PD-L1 and up-regupation of LC3-II without MHC-I changing as well (Fig.3d and Supplementary Fig.3c). The effect of calpains on PD-L1 was also been analyzed by immunofluorescence assay. The results suggested that depletion of calpains resulted in a reduction of membrane-bound PD-L1 but an augment of LC3-II as well as the co-localization between PD-L1 and LC3-II in cytoplasm. Contrastively, overexpression of calpains decreased the co-localization between PD-L1 and LC3-II but stabilized PD-L1 binding on the plasma membrane (Fig.3e and g). Comparatively, there was no co-localization between MHC-I and LC3-II regardless of the level of calpains expression (Fig.3f and h). Furthermore, we verified that amlodipine could block the calpains-mediated increase of PD-L1 proteins and triggered autophagy by co-incubation amlodipine with calpains-overexpressing tumor cells (Fig.3i-j). These findings suggest that calpains activated by intracellular calcium stabilize PD-L1 by suspending autophagy.

**Fig. 3.**
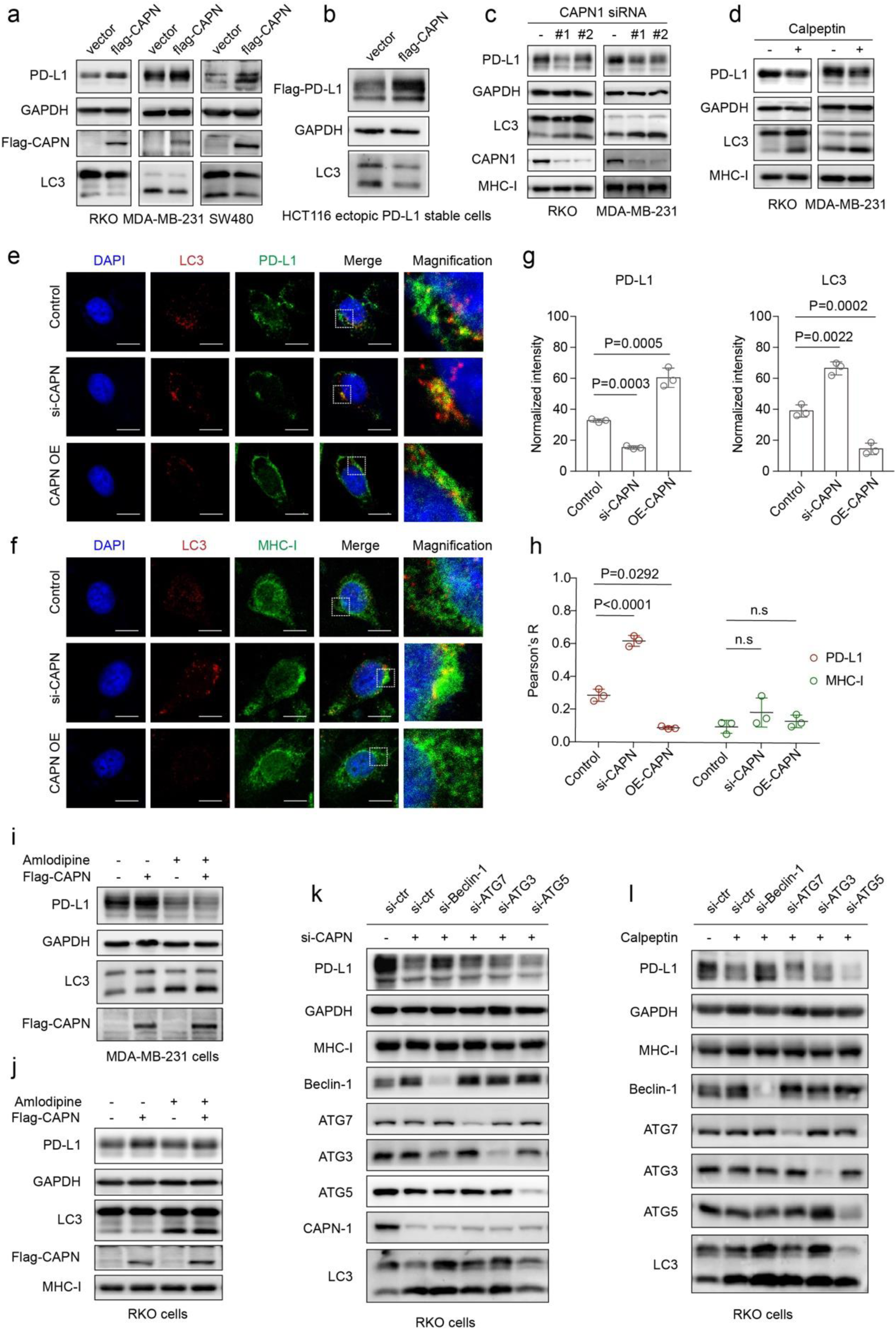
Calpains modulated the calcium-dependant stabilization of PD-L1. **a**,**b**, PD-L1 and LC3 expression analyzed by immunoblotting in the indicated cells overexpressing the Flag-tagged calpain (Flag-CAPN). **c**,**d**, the expression of PD-L1, MHC-I and LC3 detected by immunoblots in the indicated cells transfected with siRNAs for calpains or treated with calpain inhibitor calpeptin(10μM, 24h). **e**,**f**, Detection of co-localizations between LC3 and PDL1/MHC-I in RKO cells respectively transfected with siRNAs for CAPN and ectopic flag-CAPN. White-edged frames indicate the representative fields to be presented in magnification. Scale bars indicate 10μm. These experiments were respectively repeated three times independently with similar results. **g**, Quantifications of PD-L1 (left) and LC3 (right) intensity in **e**. Values are means ± s.d. from *n* = 3 independent experiments. Statistical differences were determined by ANOVA post-hoc test. **h**, Quantification of the co-localizations between LC3 and PD-L1/MHC-I in RKO cells with corresponding treatments in **i** and **j**. Values are means ± s.d. from n = 3 independent experiments. Statistical differences were determined by ANOVA post-hoc test. n.s, no significance. **i**,**j**, PD-L1, LC3 and MHC-I expression analyzed by immunoblots in the indicated cells transfected with Flag-CAPN in the presence or absence of amlodipine(25μM, 24h). **k**, Beclin-1/ATG3/ATG7/ATG5 were silenced respectively by specific siRNAs in the calpain-knockout RKO cells and detected by immunoblots with the indicated antibodies. **l**, Beclin-1/ATG3/ATG7/ATG5 were silenced respectively by specific siRNAs in the RKO cells co-incubated with calpeptin (5μM,24h), subjected to immunoblots with the indicated antibodies.

Next, we asked in the circumstance of calcium flux inhibiting, how inactive calpains contributed to the elimination of PD-L1 by inducing autophagy. We noticed that previous studies had indicated that calpains manipulated the cleavage of autophagy-related proteins, such as ATG3, ATG5, ATG7 and Beclin-1^45-48^. To find out the exact protein involved in this PD-L1 autophagy process, we silenced these candidate proteins respectively by specific siRNAs in the calpains-depleted tumor cells and found that the knockdown of calpains impaired the PD-L1 expression and increased the level of Beclin-1 and LC3-II, while the depletion of Beclin-1 was able to rescue the down-regulation of PD-L1 (Fig.3k). Similar results were acquired with the treatment of the calpain inhibitor calpepetin in the calpain-knockout tumor cells(Fig.3l). These results suggest that the effect of calcium/calpains on stabilizing PD-L1 is mediated by the autophagy repression by calpains cleavage of Beclin-1.

### Activated autophagy targeted the PD-L1 anchored on recycling endosomes under low intracellular calcium concentration

Subsequently, we investigated how calpain-dependent activated autophagy specifically targeted PD-L1. Studies showed that calpains facilitated the Rab11-marked recycling endosome to transfer and fuse to plasma membrane by recruiting Rab coupling proteins which tightly interacted with Rab11^49^, which was consist with our results shown in Fig.3e. Besides, substantial evidence indicated that the recycling endosome played an essential role in the formation of autophagosomes ^17-19 26^. Moreover, it is indicated that PD-L1 proteins sited on the plasma membrane were abundantly internalized to recycling endosome, thereby escaping from being degraded by lysosomes ^5^. Taken together, we hypothesized that by blocking calcium influx, inactivated calpains could no longer recruit recycling endosomes loaded with PD-L1 to the cell membranes, which induced the accumulation of PD-L1 on recycling endosomes and sequentially the autophagic degradation.

We identified that PD-L1 protein physically interacted and colocalized with Rab11 (the specific marker of the recycling endosome) by co-immunoprecipitation (Fig.4c) and immunofluorescence assays (Fig.4a), while MHC-I was barely co-localized with Rab11 (Fig.4b). Of note, depletion of calpains increased the co-localization between LC3-II-labelled autophagosome and Rab11-labelled recycling endosome (Fig.4d). Interestingly, the co-localizations between LC3-II and other membrane compartments where autophagy usually initiated from, such as GRP94-labelled endoplasmic reticulum (ER) (Fig.4e), ATP5A-labelled mitochondria (Fig.4f) and 58k-labelled Golgi apparatus (Fig.4g), were almost unaffected by calpains depletion (Fig.4h). Likewise, treatment with amlodipine promoted the co-localization between LC3-II-labeled autophagosome and Rab11-labeled recycling endosome but not the ER, mitochondria or Golgi apparatus (Fig.4i and Supplementary Fig.4a-d). These findings indicate that the suppression of calcium/calpains by amlodipine triggers selective autophagic degradation of RE-bound PD-L1.

**Fig. 4.**
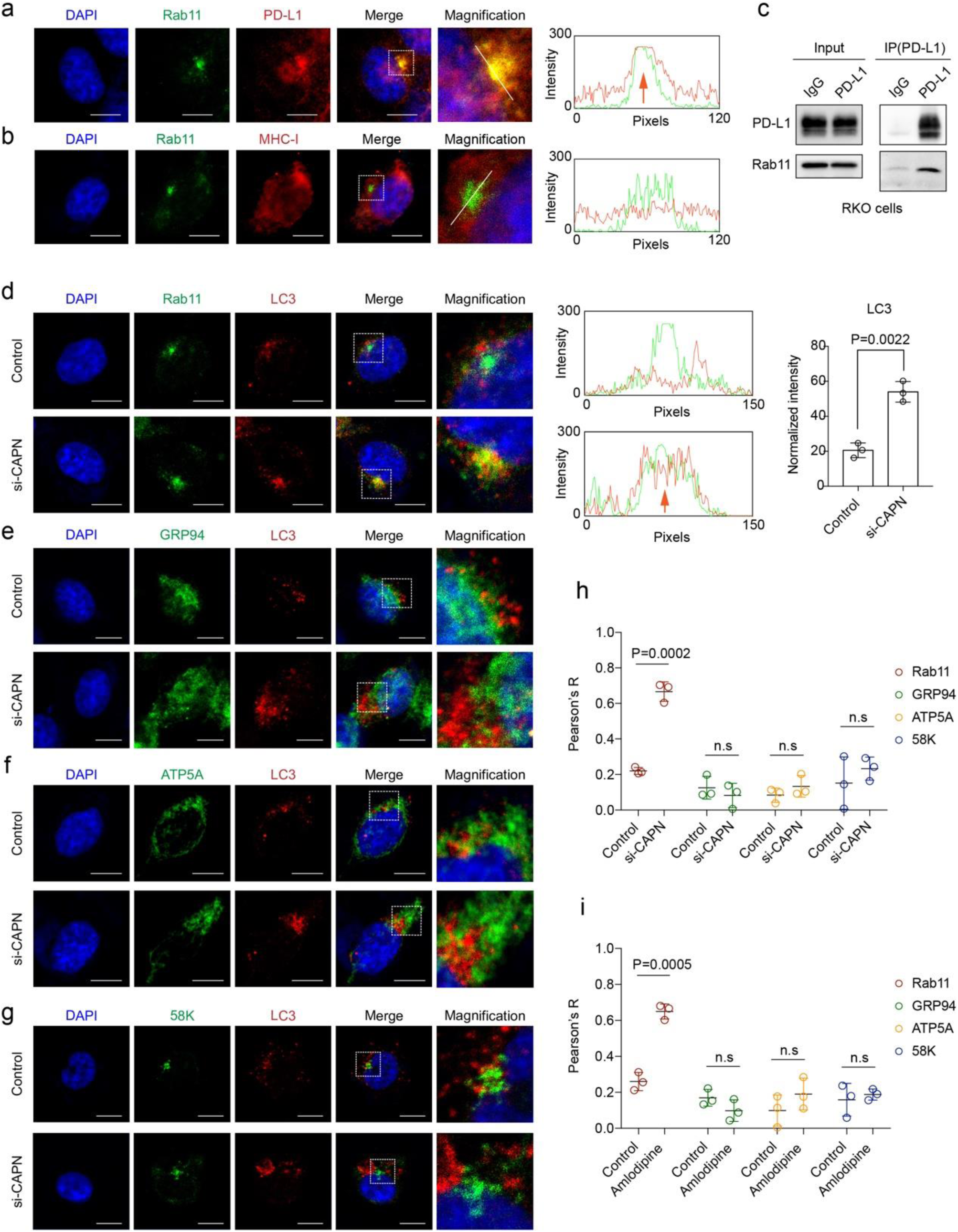
Inactivated calpains facilitated the autophagic degradation of PD-L1 from recycling endosomes. **a**,**b**, Left, immunofluorescence showing the co-localizations between endogenous Rab11 (a specific marker of recycling endosome) and PD-L1/MHC-I. Dashed frames indicate the representative areas to be shown in magnification. Scale bars indicate 10μm. These experiments were repeated three times independently with similar results. Right, the intensity profiles of PD-L1 and LC3 along the white line, with the site of co-localization indicated by orange arrows. **c**, co-IP assay showing the interaction between endogenous PD-L1 and Rab11 in RKO cells. This experiment was repeated three times independently with similar results. **d**, left, immunofluorescence showing the co-localization between LC3 and Rab11 in RKO cells transfected with siRNAs for calpains. Dashed frames indicate the representative areas to be shown in magnification. Scale bars indicate 10μm. Middle, the intensity profiles of Rab11 and LC3 along the white line. The orange arrow indicates the site of co-localization. Right, quantification of the co-localizations between Rab11 and LC3. Values are means ± s.d. from n = 3 independent experiments. Statistical differences were determined by two-sided Student’s *t*-test. **e**-**g**, RKO cells were transfected with siRNAs for calpain and detected the co-localizations between LC3 and GRP94(ER) /ATP5A(mitochondria)/58K(Golgi apparatus). Dashed frames indicate the representative areas to be presented in magnification. Scale bars indicate 10μm. **h**, Quantification of the co-localizations between LC3 and Rab11/GRP94/ATP5A/58K in RKO cells transfected with siRNAs for calpain. Values are means ± s.d. from n = 3 independent experiments. Statistical differences were determined by ANOVA post-hoc test. **i**, Quantification of the co-localizations between LC3 and Rab11/GRP94/ATP5A/58K in RKO cells with amlodipine treatment. Values are means ± s.d. from n = 3 independent experiments. Statistical differences were determined by ANOVA post-hoc test.

### Amlodipine promoted tumor-specific T cell cytotoxicity *in vitro* and *in vivo*

As amlodipine significantly decreased the expression of PD-L1 in vitro, we further explored whether it functionally affected the anti-tumor immunity. Firstly, after treatment with amlodipine, human PD-1 and Fc fragment fusion protein was incubated with tumor cells. Both immunofluorescence and FACS showed a decrease in the fluorescence intensity of PD-1 binding to the tumor cell surface (Fig.5a and b and Supplementary Fig.5a).

**Fig. 5.**
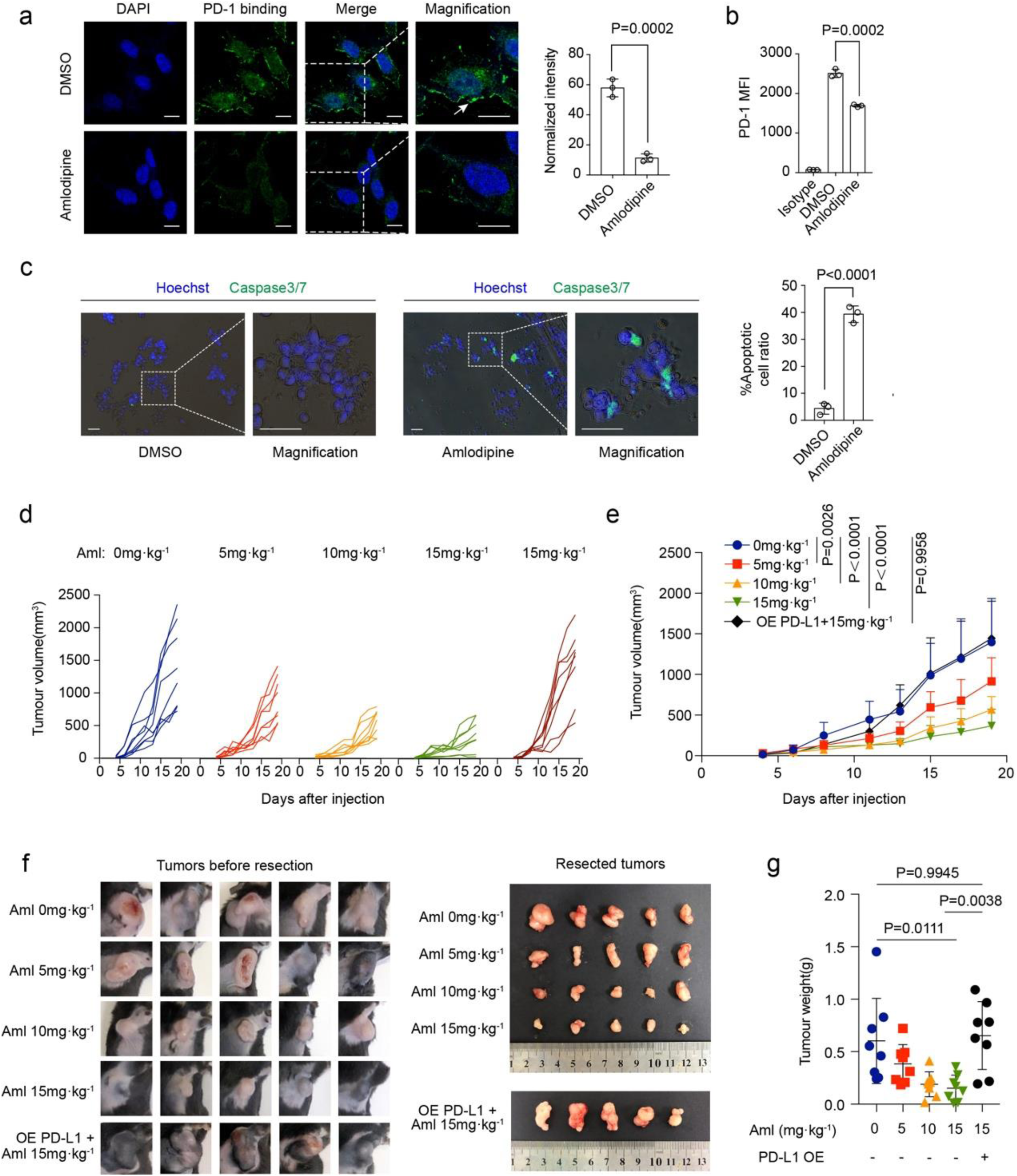
Amlodipine suppressed PD-1 binding and growth of tumor cells. **a**, Left, immunofluorescence showing the PD-1 binding on RKO cells in the presence of amlodipine. Dashed frames indicate the representatvie areas to be shown in magnification. The white arrow indicates the binding site. Scale bars, 10μm. Right, Values are means ± s.d. from n=3 independent experiments. Statistical differences were determined by two-sided Student’s *t*-test. **b**, Flow cytometry detection of PD-1 binding to RKO cells treated with amlodipine. Quantification of PD-1 mean fluorescence intensity (MFI). Values are means ± s.d. from n=3 independent experiments. Statistical differences were determined by two-sided Student’s *t*-test. **c**, T cell killing assay of RKO cells with amlodipine treatment. Cell nuclei marked by Hoechst in blue, and apoptotic cells stained by fluorescent dyes of caspase3/7 cleavage (green). The fluorescence images were merged with phase-contrast images of the cultured cells. Dashed frames indicate the representative areas to be shown in magnification. Scale bars, 50μm. Values are means ± s.d. from three independent experiments (*n* = 3). Statistical differences were determined by two-sided Student’s *t*-test. **d**, Tumor growth rate of each individual that received indicated concentrations of amlodipine after inoculation of MC38 or PD-L1 overexpressing clone (p=8 per group). **e**, The tumor growth rate of each group that received indicated concentrations of amlodipine after inoculation of MC38 or PD-L1 overexpressing clone. The points indicate means and 95% confidence intervals (two-way ANOVA; *n* = 8 per group). **f**, Left, representative tumors before resection of each group that received indicated concentrations of amlodipine sacrificed on 19^th^ day after inoculation. Right, the tumors in left after resection. **g**, Tumor weight shown as means ± s.d. (*n* = 8 per group), compared by ANOVA post-hoc test.

Then, T cell killing assay was performed by co-culturing tumor cells with activated human peripheral blood mononuclear cells (PBMCs). As a result, treatment with amlodipine contributed to activating the T-cell cytotoxicity on tumor cells and induced the apoptosis of tumor cells (Fig.5c).

Next, to investigate the anti-tumor effect of amlodipine in vivo, we established mouse colorectal MC38 subcutaneous transplanted tumor models in C57BL/6 mice. We also established an additional group of rescue condition by overexpressing mouse PD-L1 in MC38 cells. Daily peritoneal injection of amlodipine (0, 5, 10 or 15 mg·kg^−1^) began from the fourth day after inoculation of tumor cells and ended on the 19^th^ day when all groups of mice were sacrificed simultaneously. The PD-L1-overexpressing group received the same injection of amlodipine 15 mg·kg^−1^ per day. The result revealed amlodipine treatment conducted a dose-dependent inhibition of tumor growth, and the alleviation of tumor growth could be rescued in the amlodipine/PD-L1-overexpressing MC38 tumor models (Fig.5d-g and Supplementary Fig.5b). Collectively, these data suggest that amlodipine functionally suppresses PD-1 binding and growth of tumor cells, and induce the T cell toxicity, which highlight amlodipine as a promising agent for tumor immunotherapy.

### PD-L1 dependent effects of Amlodipine in promoting anti-tumor immunity

Next, we aimed to investigate whether amlodipine promoted anti-tumor efficacy by inducing autophagic degradation of PD-L1. Tumor tissues of the amlodipine-treatment group observed by immunofluorescence showed increased co-localizations between PD-L1 and LC3-II-labelled autophagosome compared with the control group, but no change in the co-localization between MHC-I and LC3-II. Consistently, increased co-localizations between LC3-II-labelled autophagosome and Rab11-labelled recycling endosome were identified in the group treated with amlodipine(Fig.6a). Furthermore, compared with the control and amlodipine/PD-L1-overexpressing group, the group that received amlodipine 15 mg·kg^−1^ per day significantly down-regulated the expression of PD-L1 detected by immunohistochemistry assay (Fig.6b) and immunoblots (Supplementary Fig.6a), and up-regulated the infiltration of CD8^+^ T cell in tumor tissues by analysis of immunohistochemistry assay (Fig.6d). Intriguingly, in myocardium tissues of the same individuals, there was no significant difference of PD-L1 expression or CD8^+^ T-cell infiltration between the control group and the amlodipine-treatment group. Besides, compared with the Rab11 expression in tumor tissues, the expression of Rab11 in myocardium tissues of the same indivisual was much lower (Fig.6c), which might expain why this calcium-associated specific regulation of RE-bound PD-L1 tends to take place in the sites with high-expression of RE like tumor tissues, but has no effect on normal tissues with low-expression of RE like myocardium tissues. Taken together, these data imply that amlodipine has potential therapeutic effect on anti-tumor immunity by decreasing PD-L1 in vivo.

**Fig. 6.**
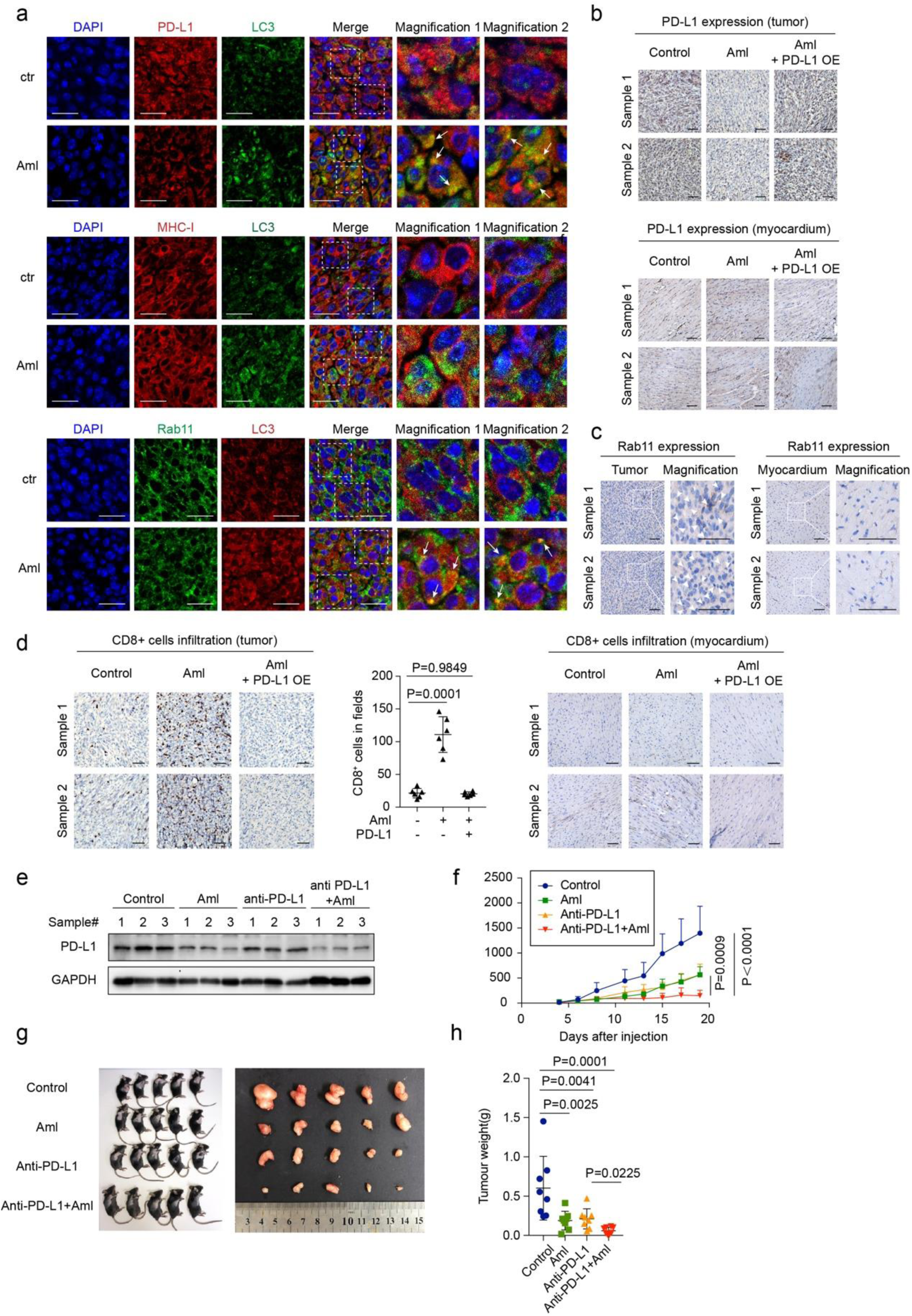
Amlodipine promoted anti-tumor immunity by decreasing PD-L1 expression in vivo. **a**, Immunofluorescence showing the colocalizations between LC3 and PD-L1/MHC-I/Rab11 in the indicated tumor tissues. Dashed frames indicate the representative areas to be shown in magnification. The white arrows indicate the co-localizing sites. Scale bars, 20μm. This experiment was conducted in four independent samples of each group with similar results. **b**, Immunohistochemistry showing PD-L1 expression in the indicated tumor tissues and the myocardium tissues of the same individuals. Scale bars, 100μm. **c**, Immunohistochemistry showing Rab11 expression in the tumor tissues and the myocardium tissues of the same individuals. Scale bars, 100μm. **d**, Immunohistochemistry showing CD8^+^ cell infiltration in the indicated tumor tissues (left) and the myocardium tissues (right) of the same individuals, and quantification of CD8^+^ cell infiltration (middle) in the tumor tissues. Scale bars, 100μm. Statistical results shown as means ± s.d. in each group (*n* = 8), determined by ANOVA post-hoc test. **e**, PD-L1 expression detected by immunoblots in the indicated tumor tissues. **f**, The tumor growth rate of each group that received indicated treatments after inoculation of MC38 tumor cells. The points indicate means and 95% confidence intervals (two-way ANOVA; *n* = 8 per group). **g**, Representative individuals of each group that received the indicated treatments sacrificed on 19^th^ day after inoculation. **h**, Tumor weight shown as means ± s.d. (*n* = 8 per group), determined by ANOVA post-hoc test.

**Fig. 7.**
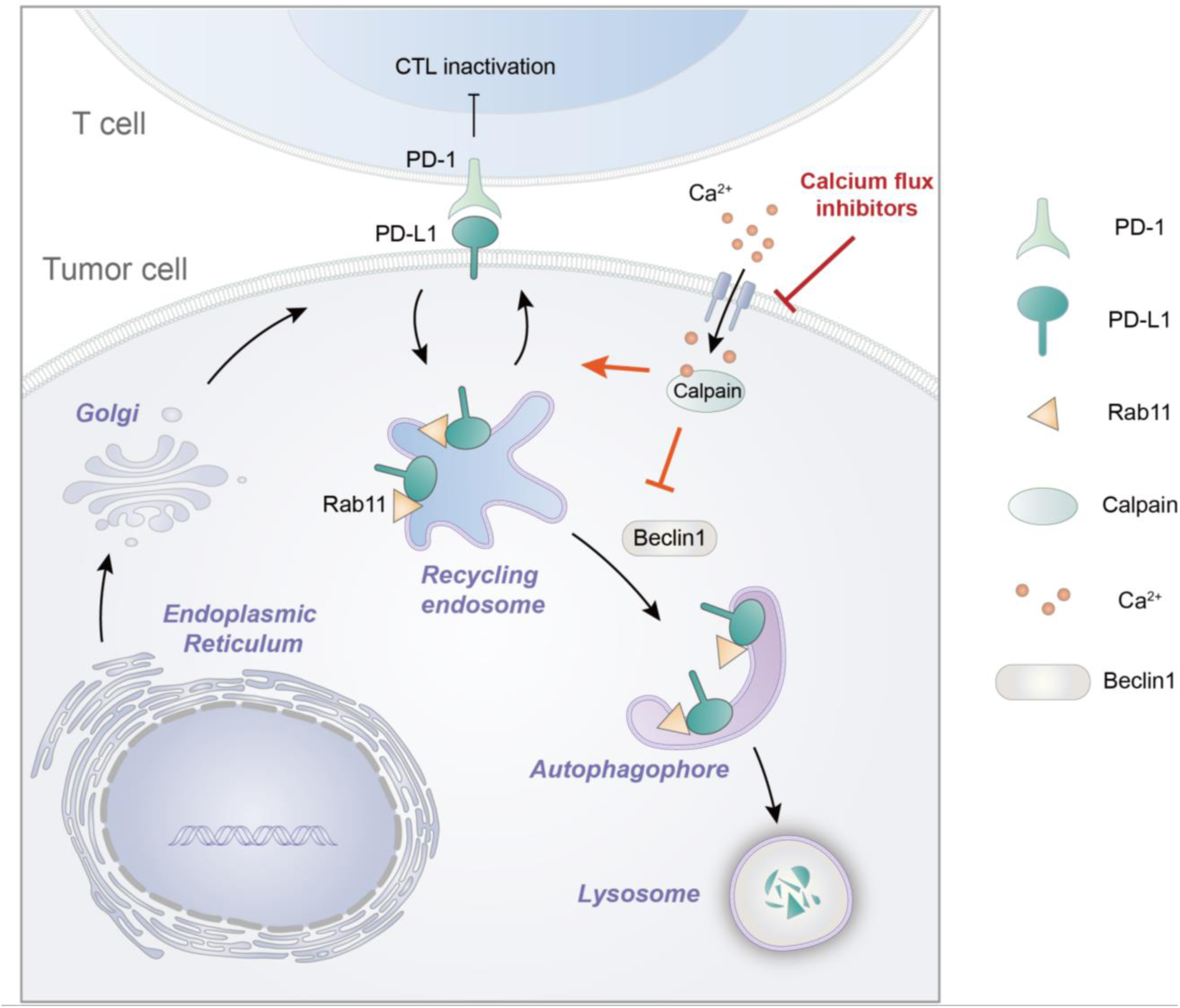
The calcium flux inhibitor Amlodipine induces PD-L1 degradation by promoting selective autophagy of recycling endosome. Calpain, when bound to calcium, inhibits the autophagy of recycling endosome (RE) by cleaving Beclin1, thereby stabilizing PD-L1 expressed on RE. The blockade of calcium flux by Amlodipine suppressed calpain function and induced PD-L1 autophagy, blocking PD-L1 dependent tumor immune evasion *in vivo*.

Besides, we set other two groups, which were treated with purified anti-mouse PD-L1 agent (5mg·kg^−1^), and a combination of amlodipine (10mg·kg^−1^) and the anti-PD-L1 agent (5mg·kg^−1^) to compare the therapeutic effect on tumor growth. Intriguingly, analysis of tumor tissues by immunoblots denoted that amlodipine treatment decreased the expression of PD-L1 while anti-PD-L1 antibody had little effect on the expression of PD-L1 (Fig.6e). Moreover, amlodipine promoted the anti-tumor effect of anti-PD-L1 antibody (Fig.6f-h and Supplementary Fig.6b and c), probably because the two therapies depended on different blocking mechanisms of PD-L1. Collectively, these findings suggest that amlodipine can effectively improve the efficacy of anti-PD-L1-dependent immunotherapy by promoting the autophagic degradation of PD-L1.

## DISCUSSION

In this work, we demonstrated that the calcium flux inhibitor amlodipine can selectively induce the autophagic degradation of PD-L1 in a calcium-dependent manner. Functionally, amlodipine not only suppresses PD-1 binding and augments T cell cytotoxicity, but also alleviates tumor growth and enhances the efficacy of the anti-PD-L1 agent in mice. As a canonical calcium channel blocker, amlodipine is widely used in treatment of lowering blood pressure, however, its function involved in ant-tumor effect has not yet been fully brought to light. A recent study proposed that amlodipine could be a potential inhibitor for selective uveal melanoma, but the exact mechanism remained unknown ^50^. Compared with nifedipine and other dihydropyridine CCBs, amlodipine owns the longest half-life which can effectively keep the plasma drug concentration with once-daily dosing^38^. These data suggest that amlodipine may be a promising adjuvant drug for anti-PD-L1 immune checkpoint blockade.

In this study, we highlight a novel role that calcium plays in PD-L1 regulation. Our data emphasize that the intracellular calcium flux positively modulates PD-L1 in the protein level rather than the mRNA level. Under high concentration of intracellular calcium, activated calcium-sensitive proteases calpains recruit recycling endosomes where a substantial proportion of PD-L1 localized on to transfer and fuse to the plasma membrane and conduct the cleavage of Beclin-1 to suspend autophagy activity. Given the fact that calcium signaling is perturbation in tumor cells ^51 52^ and several calcium channel blockers have been used as targets for cancer therapies ^52 53^, it is reasonable to speculate that aberrant high concentration of intracellular calcium in tumor cells is prone to stabilize PD-L1 and helps tumor cells evade T-cell killing. Under effect of amlodipine, calpains turn inactivated by the decline of intracellular calcium, which blocks the RE-bound PD-L1 trafficking to cell membranes, and exempts Beclin-1 from been cleaved. Ultimately, accumulated PD-L1 on the recycling endosomes are degraded by activated RE-associated autophagy.

Generally, autophagy is considered as a nonspecific catabolic process that sustains the cellular homeostasis. Increasing studies have proposed the cases of selective autophagy, in which the phagophores can specifically recognize and engulf the targets. It has been suggested that L-type calcium channel blockers enhance autophagy by regulating the activity of the calcium-sensitive proteases calpains rather than a mTOR-dependent manner ^43^. Calpains may cleave several autophagy-related Atg proteins including Beclin-1, ATG3, ATG5, and ATG7 ^45-48^. Beclin-1 (the mammalian homolog of Atg6) is a component of type III phosphoinositide 3-kinase (PI3-K) complex, playing a key role in the development of autophagosome and tumor suppression ^54 55^. Nonetheless, the calpain-regulated autophagy has not been linked to the selectivity of autophagy towards various membrane structures. Particularly, our data propose a specific calcium-dependent selective autophagy of RE-bound PD-L1, which does not imperil other membrane proteins like MHC-I, or involving other autophagy-associated membrane compartments such as ER, Golgi apparatus or mitochondria. Besides, compared with other normal tissuses with low expression of RE like myocardium tissues, the tumor tissues with high-expression of RE are more likely to response to this calcium-dependent degradation of RE-bound PD-L1.

To date, prevalent anti-PD-L1 agents take effect by targeting PD-L1 on tumor cell surface. Nevertheless, these drugs do not cope well with the compensatory redistribution or intracellular PD-L1. And a large proportion of patients do not show durable responses or get acquired resistance after long-term therapy. In this regard, we exploited the calcium flux inhibitor, which facilitate the degradation of PD-L1 stored on recycling endosomes and thus eliminate PD-L1 from the inside, as a complementary therapeutic strategy for anti-PD-L1 antibody. The in-vivo experiments show that the combined treatment of anti-PDL1 antibody and amlodipine can achieve a synergistic effect and enhance the efficacy of immunotherapy of tumor.

In summary, our study sheds light on a potential effect of amlodipine on PD-L1 suppression through a calcium-dependent pathway, and a novel selective degradation of autophagy initiated from recycling endosome, which provides a new therapeutic strategy for immunotherapy.

## METHODS

### KEY RESOURCES TABLE

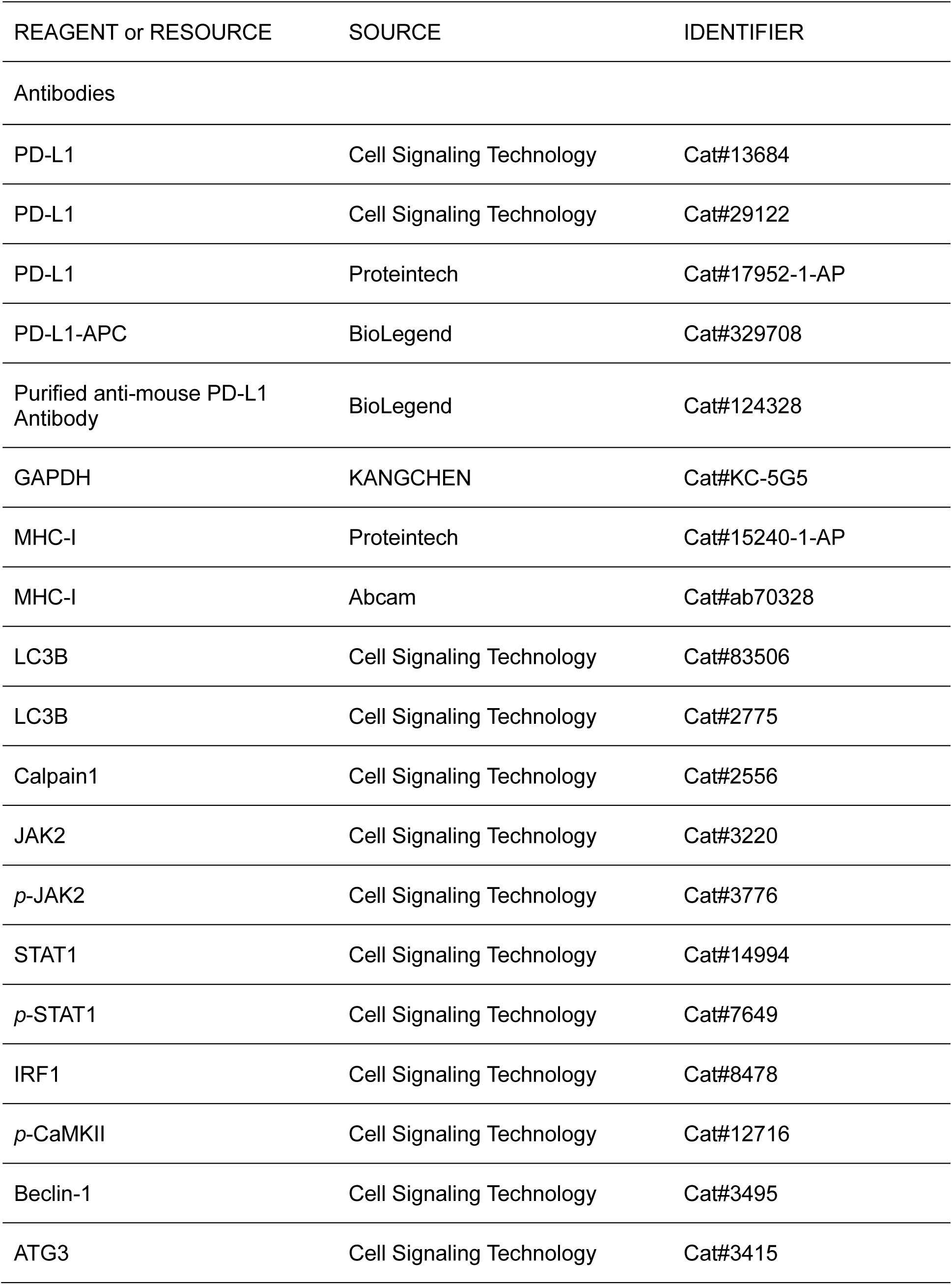
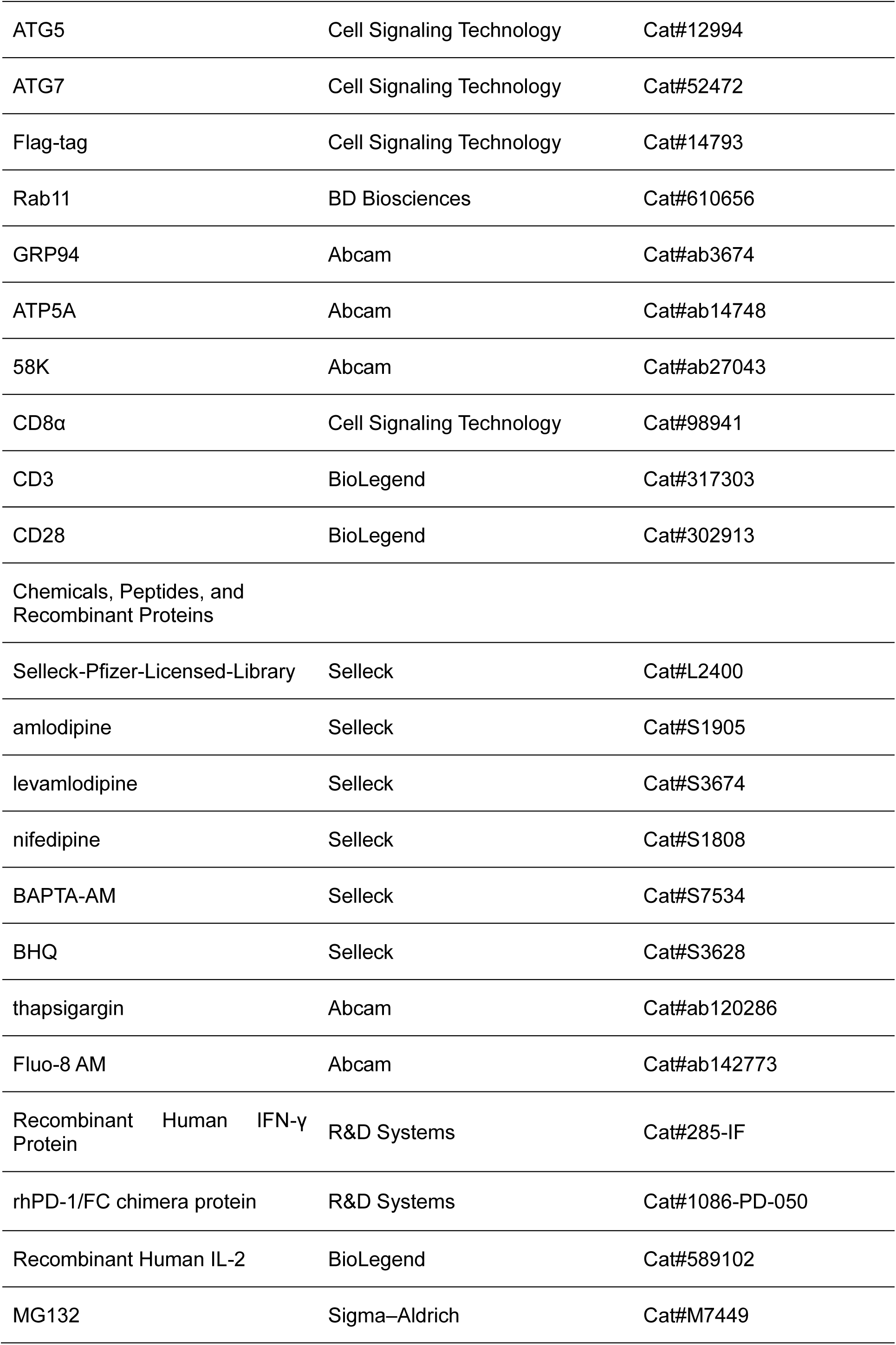
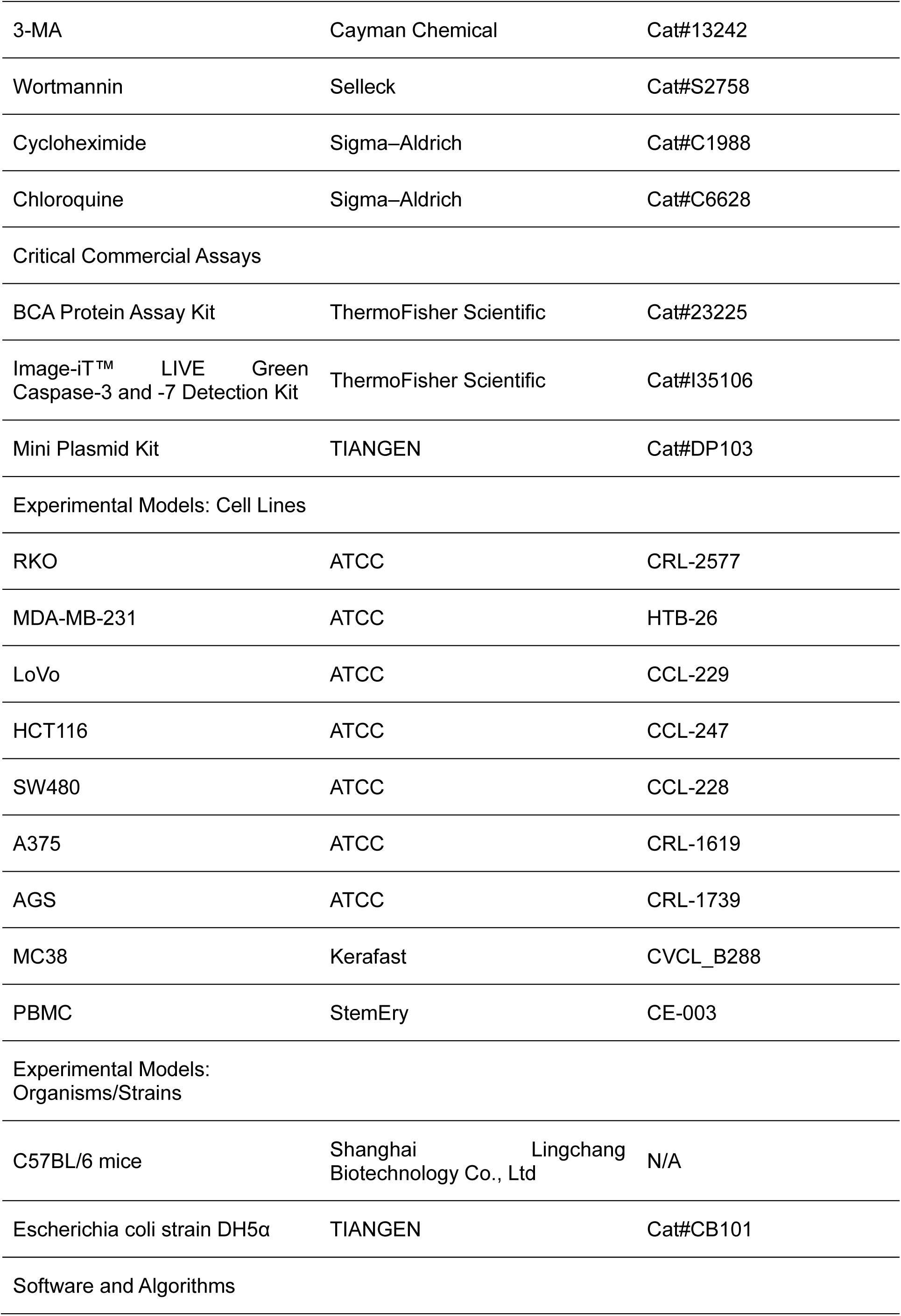
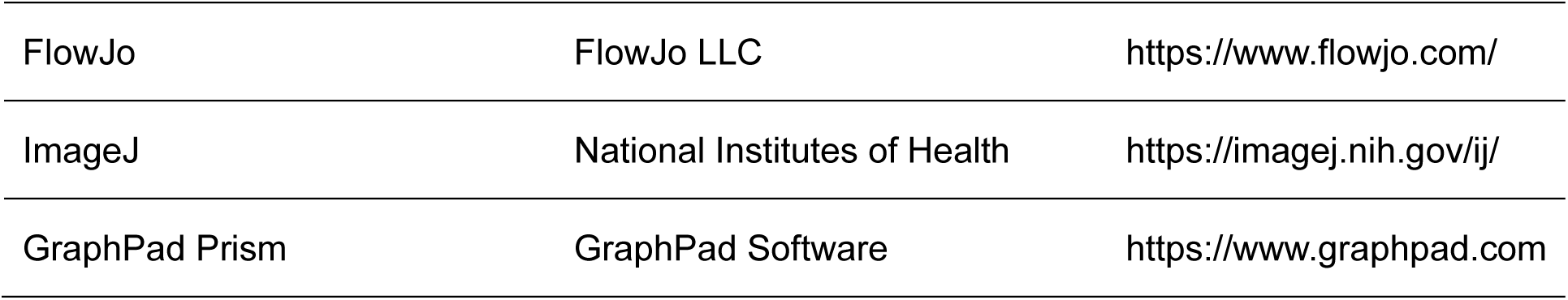

### CONTACT FOR REAGENT AND RESOURCE SHARING

For further details of reagents and resources, please contact the Lead Contact, Jie Xu.

### EXPERIMENTAL MODEL AND SUBJECT DETAILS

All animal experiments were executed in accordance with the ethical obligations approved by the department of laboratory animal science of Fudan University and the Institutional Animal Care and Use Committee of Renji Hospital, School of Medicine, Shanghai Jiaotong University.

All the C57BL/6 mice (4 weeks old, female) were randomly numbered and divided into different groups for further treatment (n=8 for each group). After one-week adaption of the environment, each group of mouse received subcutaneous injection of mouse colorectal MC38 cells (1.5×10^6^ cells), except one group received mouse-PD-L1-overexpressed MC38 cells (1.5×10^6^ cells) as a rescue condition (the establishment of stable clones as described later). Peritoneal injection of amlodipine at different concentrations (0, 5, 10 or 15 mg•kg^−1^, q.d) were respectively given to four groups from the fourth day after inoculation, lasting about three weeks or till the end point. The PD-L1-overexpressing group received the same injection of amlodipine 15 mg·kg^−1^ per day. To compare the therapeutic effect of amlodipine and the anti-PD-L1 agent on tumor growth, other two groups were respectively treated with a purified anti-mouse PD-L1 agent (BioLegend cat#124328) (5mg·kg^−1^, i.p, at days 4, 10, and 16 after inoculation) and a combination of amlodipine (10mg·kg^−1^, i.p, q.d) and the anti-PD-L1 agent (5mg·kg^−1^, i.p, at days 4, 10, and 16 after inoculation). All animal experiments were performed in the department of laboratory animal science of Fudan University.

The tumor volume of each mouse was evaluated every 2-3 days by measuring the length and width and calculated according to the formula 1/2 × A × B^2^ (A and B indicate the long and short axis of the tumor, respectively). In concert with the ethical standards, mice were sacrificed once the tumor volume exceeded 2cm^3^ or ulcers occurred. All the mice were sacrificed on the 19^th^ day after inoculation. After the sacrifice of mice, the tumors were weighed and then mechanically disaggregated and pre-treated for further experiments.

## METHOD DETAILS

### Cell culture

All cell lines involving human colorectal cancer RKO, LoVo, HCT116, SW480 cells, human breast cancer MDA-MB-231 cells, melanoma A375 cell and mouse colorectal cancer MC38 cells were purchased from ATCC, detected negative of mycoplasma. RKO, LoVo, SW480 cells were cultured in RPMI-1640 (Gibco), supplemented with 10% FBS (Invitrogen). HCT-116 cells were cultured in McCoy’s 5A (Gibco), supplemented with 10% FBS (Invitrogen). MDA-MB-231 cells, A375 and MC38 cells were cultured in DMEM (Gibco), supplemented with 10% FBS (Invitrogen). All the cell lines were incubated at 37 °C under 5% CO_2_.

### Plasmids construction

The pcDNA3.1-Flag-PD-L1 vector was constructed by inserting synthesized cDNA encoding 3 × Flag tag and PD-L1 (clone R8443-2, Generay Biotech) into pcDNA3.1 vector using EcoRI/XhoI MCS.

The expression vectors encoding pcDNA3.1-FLAG-CAPN1 (human) was generated by inserting synthesized cDNA encoding 3×FLAG tag and calpain1 (52017-11, GENECHEM) into the pcDNA3.1 vector, using the EcoRI/XhoI multiple cloning sites (MCS). All vectors were confirmed by sequence analysis and immunoblotting with specific antibodies in which the molecular weights of the proteins were consistent with the predictions.

### Transfection of plasmids or siRNAs

For transfection of plasmids, 24h after 40% confluent cells were seeded and attached, we transfected the plasmid with the system containing 1.5μg plasmid, 4.5μl FuGENE HD (Promega) and 100μl Opti-MEM per well according to the manufacturer’s instructions, with a blank vector as the negative control. The specific siRNAs purchased from Genepharma were for the knockdown of capain1, Beclin-1, ATG3, ATG5, ATG7. DharmaFECT 1 transfection reagent (Invitrogen) was used for the transfection of siRNAs according to the manufacturer’s instructions, and the nonspecific siRNA was used as the negative control.

### Establishment of PD-L1 stable cells

We chose HCT116 cells to establish a PD-L1 stable cell line for its low expression of endogenous PD-L1. Similar to the transfection of other plasmids, we transfected ectopic FLAG-tagged PD-L1 to the HCT116 cells with the system of 1.5μg plasmid, 4.5μl FuGENE HD(Promega) and 100μl Opti-MEM per well, with a blank vector control and an empty control. After approximately two-week culture with the medium containing 800 μg/ml G418 (Gibco BRL) and the removal of dead cells, the discrete colonies formed and were identified by immunoblots. Ultimately, we chose a colon which expressed highest level of PD-L1 and generated the HCT116 PD-L1 stable cells.

### Lentivirus transduction

To overexpress mouse PD-L1 in MC38 cells, the lentiviral vector LV-Cd274-EGFP was constructed and transfected at MOI of 150 in DMEM complete medium containing 5 mg/ml polybrene. Viruses loaded with blank vector were used as a control. After twenty-four-hour culture, the medium was replaced by fresh DMEM complete medium for the removal of inactive viruses and dead cells. The cells were incubated for another 72h before the identification of transfection efficiency by fluorescent microscopy. The PD-L1 expression were then tested by qPCR and immunoblots.

### Quantitative real-time PCR assays

The treated cells were lysed with Trizol reagent (Invitrogen) after washed twice with PBS. The extracted total RNA was reverse transcribed using the PrimeScript RT Reagent Kit (Takara) according to the manufacturer’s instructions in a 20μL reaction system. Quantitative real-time PCR was performed using an Applied Biosystem 7900HT Sequence Detection System (Applied Biosystems) with GoTaq RT-qPCR system (Promega). The Ct values were analyzed using the 2^-ΔΔCt^ method and the final results were presented as relative fold change. The expression of GAPDH served as internal reference. Each the qPCR assay was repeated three times from cell culture.

### Immunoblotting and co-immunoprecipitation

Cells were treated with RIPA lysis buffer (Beyotime) containing 1% cocktail of proteinase and phosphatase inhibitor and PMSF (ThermoFisher Scientific) after washed twice with PBS. The cell lysates were collected and centrifuged (12000rpm, 15min, 4°C). After detected the protein concentration using BCA Protein Assay Kit (ThermoFisher Scientific), the supernatant was added 1×SDS-PAGE sample loading buffer (Beyotime) and heated 95°C for 10 min. The protein samples were loaded in appropriate concentrations of SDS–PAGE for electrophoresis, and subsequently transferred to PVDF membranes (Bio-Rad). After blockade with 5% bovine serum albumin (ThermoFisher Scientific) in 1×TBS buffer (Sangon Biotech) for one hour, the membranes were incubated with the primary antibodies at 4°C overnight. The membranes were washed five times with TBST buffer (1×TBS, 0.1%Tween20) before incubated with the secondary antibodies labelled with HRP (KANGCHEN) at room temperature for one hour. The membranes were washed five times with TBST and detected by ChemiDoc imaging system (Bio-Rad).

For co-immunoprecipitation, cells were lysed with IP Lysis Buffer (ThermoFisher Scientific) containing 1% cocktail of proteinase and phosphatase inhibitor and PMSF (ThermoFisher Scientific). The cell lysates were collected and centrifuged (15000rpm, 2min, 4°C). The supernatant was collected and treated with DNase (QIAGEN) for 20 min at room temperature. Each sample (300μL) was taken 16μL lysate added 4μL 5×SDS-PAGE sample loading buffer (Beyotime) as the inputs, while the rest was incubated with 1μg primary antibody or IgG as controls, followed by slow-speed rotation at 4 °C overnight. Meanwhile, Protein G Agarose beads (20398, ThermoFisher Scientific) blocked with 4% BSA in TBS were also rotated at 4 °C overnight. The blocked beads were then added equally to each sample, followed by another slow-speed rotation at room temperature for one hour. After washed four times with PBS with high-speed rotation, samples were eluted in 30μL SDS sample buffer at 95 °C for 5 min. The inputs and the immunoprecipiated samples were subjected to immunoblotting as described above.

### Immunofluorescence

Cells seeded in 8-well chamber slides (C7182, Sigma) were removed the culture medium, washed twice with PBS and then fixed with 4% formaldehyde (28908, Thermo Fisher) for 20 min. After washed twice with PBS, cells were permeabilized and blocked with 0.2% Triton X-100 and 1% BSA in PBS simultaneously at room temperature for one hour. Then cells were incubated with primary antibodies at 4°C overnight. After PBS washing for five times, cells were incubated with secondary antibodies at room temperature for twenty minutes, followed by PBS washing for three times, protected from light. After nuclear staining using DAPI (#0100-20, SouthernBiotech), the slides treated with Prolong Gold reagent were observed using Zeiss LSM710 confocal microscope (Carl Zeiss). The quantification of fluorescence intensity and the colocalization analysis was evaluated using ImageJ software.

### Analysis of PD-L1 degradation rate

After pre-treatment of amlodipine with or without the indicated small molecular inhibitors, cells were incubated with cycloheximide (C104450, Sigma) (50μg/mL) at the indicated time points, and detected by western blot assay. The bands were quantified by ImageJ (version 2.0.0-rc-69/1.52p). Each band was selected by the same-sized rectangular frame and the intensity of the bands were measured after the subtraction of background. The intensity of each lane of PD-L1 was divided by the corresponding intensity of GAPDH for normalization. Ultimately, divided by the intensity at time point “0”, the relative intensity was acquired to present the trend.

### Identification of intracellular calcium

After treatment, cells were stained with Fluo-8 AM (ab142773) (100μl, 4μM) in HHBS at 37 °C under 5% CO2 for 1 hour, protected from light. Then after washed twice with HHBS, cells were observed with a fluorescence microscope.

### PD-L1/PD-1–binding assay by FACS

Cells were digested with trypsin and washed twice with PBS. The precipitates were collected and then incubated with 5μg/mL recombinant human PD-1 FC chimera protein (#1086-PD-050, R&D Systems) in staining buffer at room temperature for about one hour. After washed twice with staining buffer, cells were incubated with anti-human Alexa Fluor 488 dye conjugated antibody (#A-11013, ThermoFisher Scientific) at room temperature for 30 minutes, washed twice. Ultimately, cells in each tube were suspended with 300μL staining buffer and subjected to FACS in the FITC channel. The process from the incubation with secondary antibody should be protected from light. The results were evaluated using FlowJo software.

### T-cell killing assay

RKO cells were treated with amlodipine for 24h and then seeded in a 96-well plate with 1×10^6^ cells in each well. Meanwhile, human peripheral blood mononuclear cells (PBMC; #CE-003, StemEry) were stimulated with 100ng/mL CD3 antibody (#317303, BioLegend), 100ng/mL CD28 antibody (#302913, BioLegend) and 10 ng/mL IL2 (#589102, BioLegend) in RPMI-1640 complete medium for 24h. the pre-treated RKO cells were then co-cultured with the activated PBMCs at 1:10 ratio at 37°C under 5%CO2 for about eight hours. The cells were co-cultured in RPMI-1640 complete medium added fluorescence caspase-3/7 substrate (I35106, ThermoFisher Scientific) for 60 minutes and subsequently Hoechst for 5 minutes. After washed twice, the cells were observed by fluorescence microscopy using appropriate bandpass filters.

### Immunohistochemistry

The mouse MC38 tumor specimens were incubated with antibodies against PD-L1 (Cat#17952-1-AP, 1:150; Proteintech) and CD8α (Cat#98941, 1:400; Cell Signaling Technology). After washed three times with PBS, the tissues were then incubated with a biotin-conjugated secondary antibody, followed by avidin–biotin–peroxidase complex. Visualization was performed using amino-ethylcarbazole chromogen.

### Statistical analysis

Data of bar graphs denoted mean ± s.d. fold change relative to the control group of three independent experiments. GraphPad Prism 7.0a was used for graph drawing and statistical analysis. Two-sided Student’s *t*-test was used to compare the independent samples between two groups and one-way ANOVA with post-hoc test (Tukey’s or Dunnett’s test) was used when more than two groups. ImageJ (version 2.0.0-rc-69/1.52p) was used to quantified the immunofluorescence results. Representative immunofluorescence results were derived from three independent experiments. Three images from each replicate were used to evaluated the average fluorescence intensity of each replicate. The ultimate fluorescence intensity indicated mean ± s.d. of three replicates. Pearson’s correlation coefficient (R value) was used to analyze the colocalization between two proteins, indicating mean ± s.d. of three replicates, which was also based on the mean of three images of each replicate. P value below 0.05 was considered of statistical significance.

## Acknowledgements

This work was supported by National Natural Science Foundation of China (No: 81874050, 81572326, 81322036, 81421001, 81530072, 81830081 and 81772506), National Key R & D

Program of China (2016YFC0906002, 2016YFC0906002), and Startup Research Funding of Fudan University.

## Author contributions

CL, HY and HW performed experiments and analyzed data. JYF provided supports on study resources. CL and JX wrote the paper. JX conceived the study.

