## Supplementary Information for "Repurposing screen identifies Amlodipine as an inducer of PD-L1 degradation and antitumor immunity"

### Supplementary figures

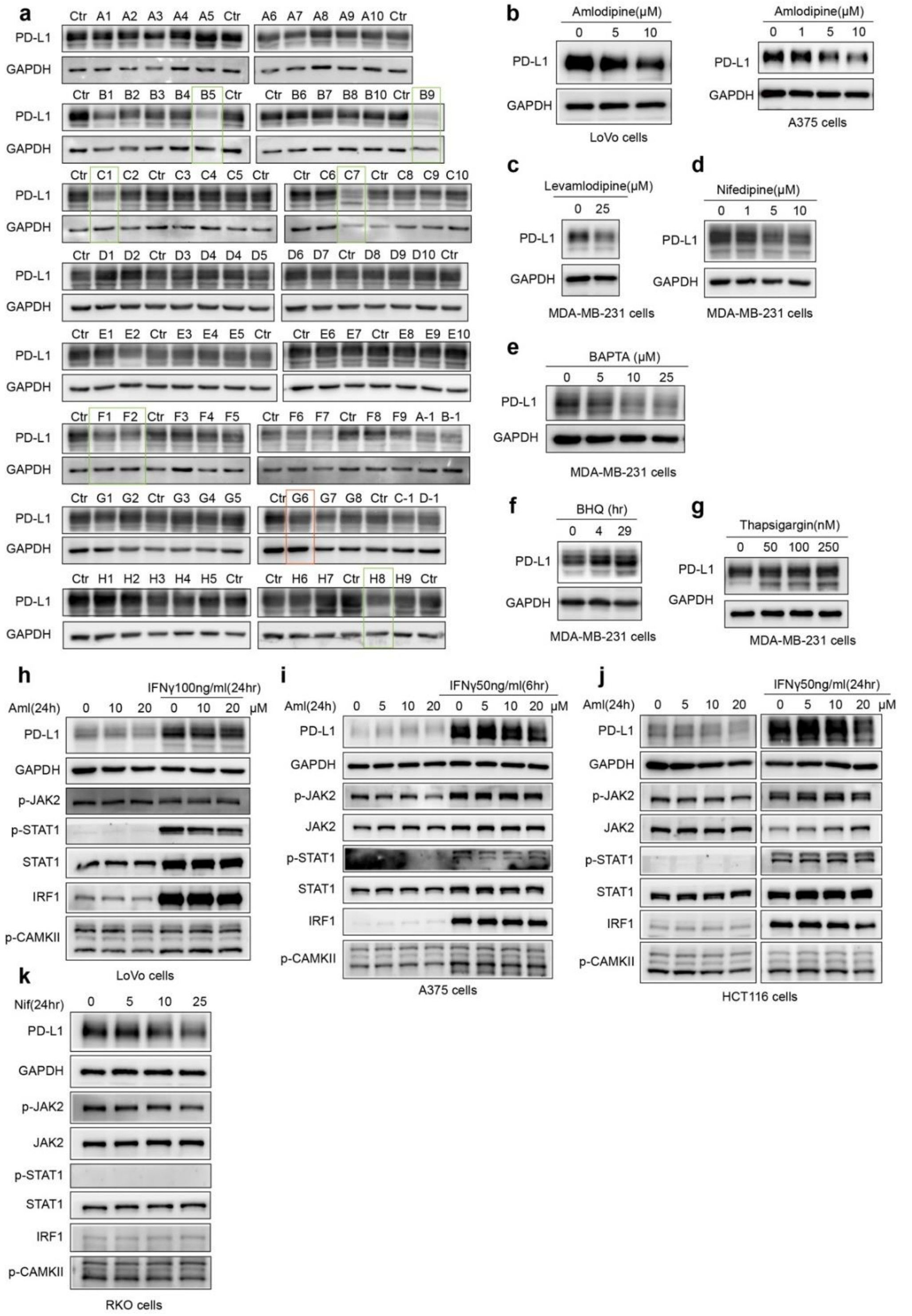


**Supplementary Fig.1 Amlodipine decreased PD-L1 expression post-translationally by blocking intracellular calcium flux. a,** The compound library screening for potential PD-L1 inhibitors. RKO cells treated with the inhibitors respectively from the clinical compound library (Selleck-Pfizer-Licensed-Library) (10μM, 24h), subjected to western blot with PD-L1 and GAPDH antibodies. Green-edged frames indicate the potential inhibitors of PD-L1 that have been reported for their correlation with PD-L1. The Orange-edged frame indicates the potential inhibitor of PD-L1 which have not been reported yet. The list of the inhibitors and their information provided in Supplementary Table 1. **b**, Incubation with amlodipine at the indicated concentrations in the indicated cells, detected by western blot. This experiment was repeated twice independently with similar results. **c**, MDA-MB-231 cells treated with Leamlodipine at the indicated concentrations for 4h. **d**, MDA-MB-231 cells treated with nifedipine at the indicated concentrations for 24h. **e**, MDA-MB-231 cells treated with BAPTA at the indicated concentrations for 3h. **f**, MDA-MB-231 cells treated with BHQ (50μM) for 0-29h. **g**, MDA-MB-231 cells treated with thapsigargin at the indicated concentrations for 4 h. **c**-**g** were repeated three times independently with similar results. **h**-**j**, LoVo, A375 and HCT116 cells treated with amlodipine(Aml) and IFN-γ at the indicated concentrations for 24h, subjected to immunoblotting with PD-L1, GAPDH, p-JAK2, JAK2, p-STAT1, STAT1, IRF1 and p-CAMKII antibodies. **k**, RKO cells treated with nifedipine(Nif) at the indicated concentrations for 24h, detected by western blot with the indicated antibodies.

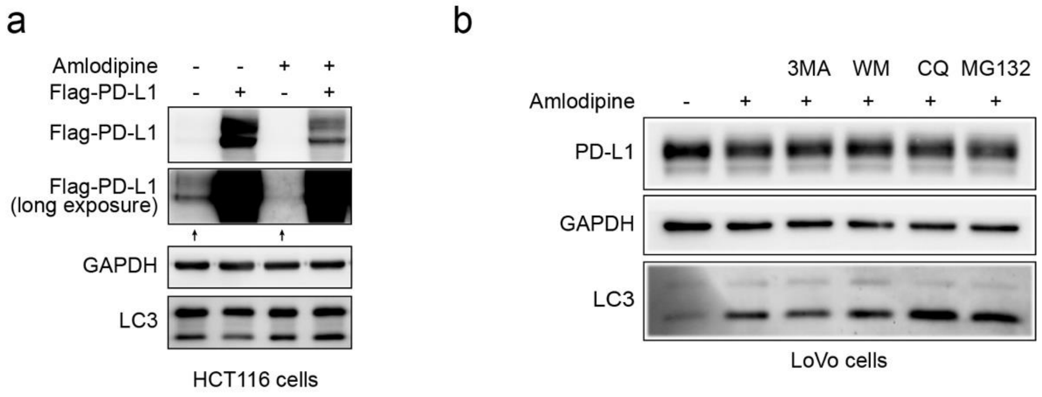


**Supplementary Fig.2 Amlodipine facilitated the autophagic degradation of PD-L1. a**, Immunoblots showing the effect of amlodipine on PD-L1, LC3 and MHC-I expression in HCT116 cells transfected with ectopic PD-L1. **b**, Immunoblots showing the effect of amlodipine (20μM, 24h) on PD-L1 and LC3 expression in the absence or presence of autophagic inhibitors 3-Methyladenine (3-MA) and wortmannin(WM), lysosomal inhibitor chloroquine (CQ) or proteasomal inhibitor MG-132.

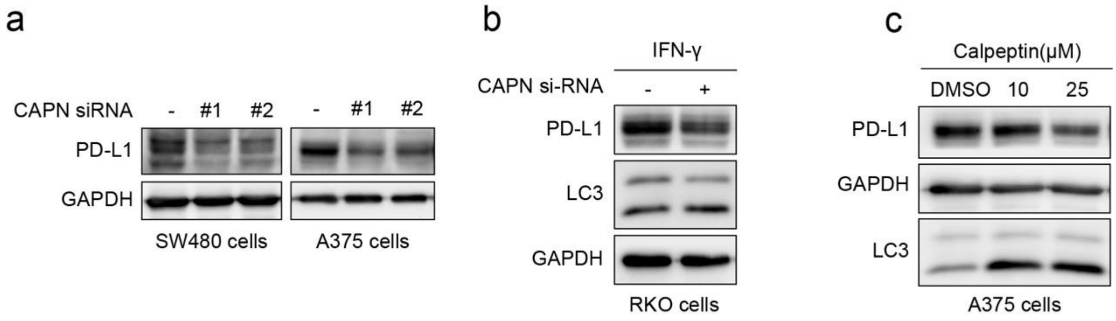


**Supplementary Fig. 3 Calpains modulated the calcium-dependant stabilization of PD-L1. a**, The expression of PD-L1 detected by immunoblots in the indicated cells transfected with siRNAs for calpains. **b**, Immunoblotting showing PD-L1 and LC3 expression in RKO cells transfected with siRNAs for calpains in the precence of IFNγ(100ng/mL, 24h). **c**, The expression of PD-L1 and LC3 detected by immunoblots in A375 cells treated with calpain inhibitor calpeptin at the indicated concentration for 4h.

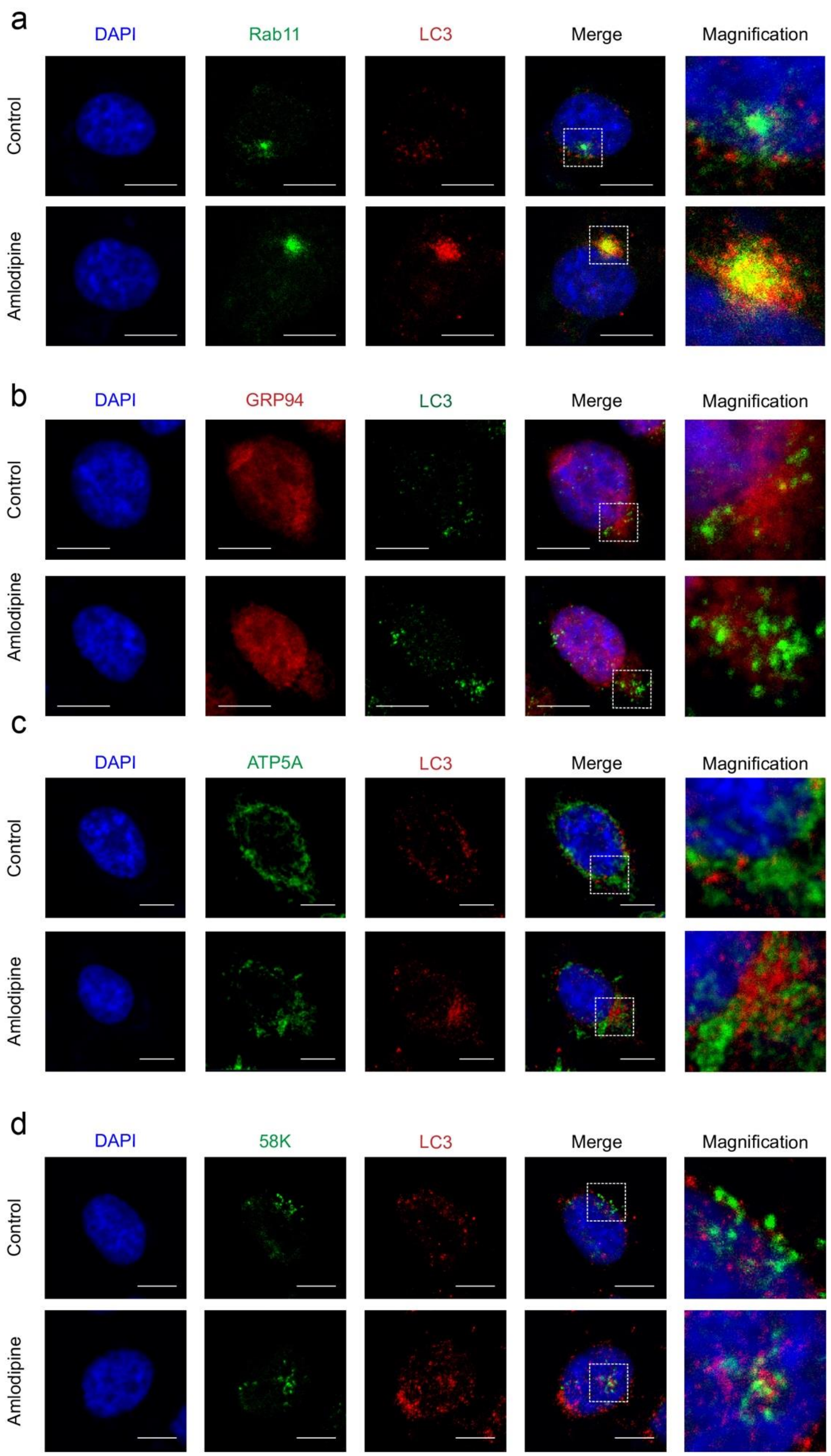


**Supplementary Fig.4 Inactivated calpains facilitated the autophagic degradation of PD-L1 from recycling endosomes. a-d**, Detection of the colocalizations between LC3 and Rab11(recycling endosome)/GRP94(ER) /ATP5A(mitochondria)/58K(Golgi apparatus) in RKO cells treated with amlodipine. Dashed frames indicate the typical areas to be presented in magnification. Scale bars indicate 10μm. These experiments were respectively repeated three times independently with similar results.

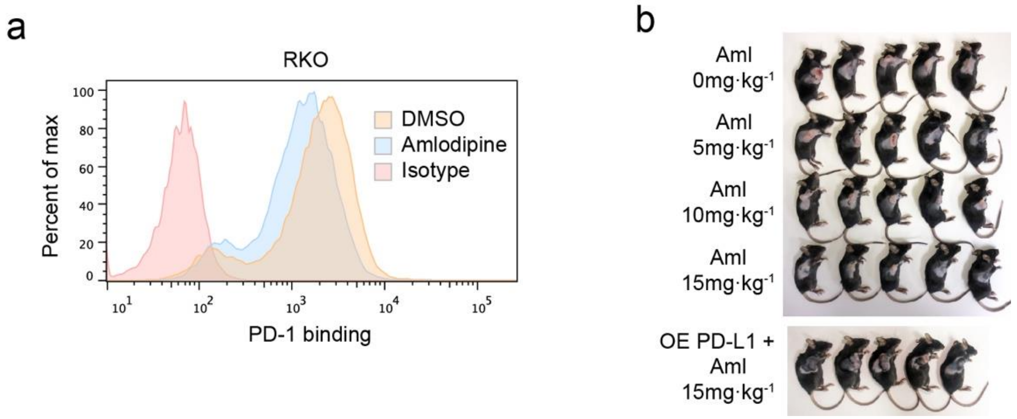


**Supplementary Fig.5 Amlodipine suppressed PD-1 binding and growth of tumor cells.** **a**, flow cytometry detection of PD-1 binding to RKO cells treated with amlodipine. **b**, Representative individuals of the indicated groups sacrificed on 19th day after inoculation.

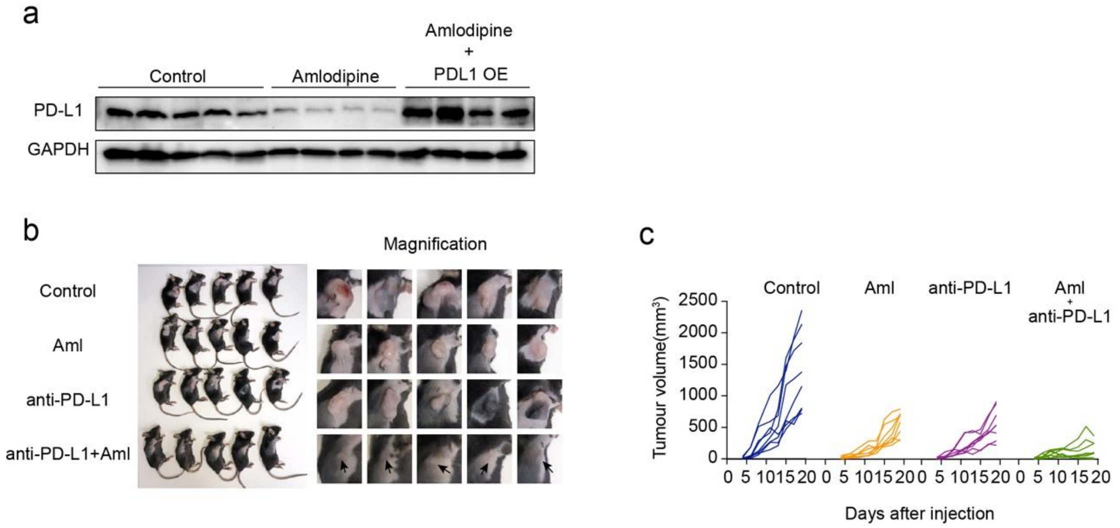


**Supplementary Fig.6 Amlodipine promoted anti-tumor immunity by decreasing PD-L1 expression in vivo. a**, PD-L1 expression detected by immunoblots in tumor tissues received indicated treatments. **b**, Left, representative individuals of the indicated groups sacrificed on 19$^{th}$ day after inoculation. Right, the corresponding magnified images of tumors. **c**, tumor growth rate of each individual that received indicated treatments after inoculation of MC38 (p=8 per group).

**Supplementary tables**

**Supplementary table 1. The information of the compound library screening for potential PD-L1 inhibitors.** (**Selleck-Pfizer-Licensed-Library, L2400).**

| *No.* | **Product Name** | **Information** |
|---|---|---|
| *a1* | Axitinib | inhibitor of VEGFR1, VEGFR2, VEGFR3, PDGFRβ and c-Kit |
| *a2* | CP-724714 | inhibitor of HER2/ErbB2, selectivity against EGFR, InsR, IRG-1R, PDGFR, VEGFR2, Abl, Src, c-Met etc in cell-free assays. Phase 2. |
| *a3* | PD173074 | FGFR1 inhibitor |
| *a4* | Lansoprazole | proton-pump inhibitor (PPI) |
| *a5* | CP-673451 | inhibitor of PDGFRα/β |
| *a6* | Glyburide (Glibenclamide) | blocker of vascular ATP-sensitive K+ channels (KATP) |
| *a7* | Amiodarone HCl | a sodium/potassium-ATPase inhibitor and an autophagy activator |
| *a8* | UK 383367 | procollagen C-proteinase inhibitor |
| *a9* | PF-3845 | FAAH inhibitor |
| *a10* | Tofacitinib (CP-690550,Tasocitinib) | a inhibitor of JAK3 |
| *b1* | Bosutinib (SKI-606) | dual Src/Abl inhibitor |
| *b2* | PD98059 | a non-ATP competitive MEK inhibitor, specifically inhibits MEK-1-mediated activation of MAPK does not directly inhibit ERK1 or ERK2. |

| | | |
|---|---|---|
| *b3* | Asenapine maleate | a high-affinity antagonist of serotonin, norepinephrine, dopamine and histamine receptors, used for the treatment of schizophrenia and acute mania associated with bipolar disorder. |
| *b4* | Tigecycline | bacteriostatic and protein synthesis inhibitor |
| *b5* | PD318088 | non-ATP competitive allosteric MEK1/2 inhibitor |
| *b6* | Levonorgestrel | female hormone that prevents ovulation. |
| *b7* | Maraviroc | CCR5 antagonist for MIP-1α, MIP-1β and RANTES |
| *b8* | Clindamycin HCl | inhibits protein synthesis |
| *b9* | PF-00562271 | ATP-competitive, reversible inhibitor of FAK |
| *b10* | Torcetrapib | CETP inhibitor |
| *c1* | PD0325901 | selective and non ATP-competitive MEK inhibitor |
| *c2* | Exemestane | aromatase inhibitor |
| *c3* | Benazepril HCl | angiotensin I converting enzyme inhibitor |
| *c4* | Linezolid | a synthetic antibiotic used for the treatment of serious infections. |
| *c5* | Sulfasalazine | a sulfa derivative of mesalazine |
| *c6* | Gemfibrozil | an activator of PPARα |
| *c7* | Atorvastatin Calcium | an inhibitor of HMG-CoA reductase |
| *c8* | Tiotropium Bromide hydrate | monohydrate of tiotropium bromide (Spiriva Tiova BA 679BR tiopropium) |
| *c9* | PF-2545920 | PDE10A inhibitor |
| *c10* | PF-4981517 | selective inhibitor of CYP3A4 (P450) |

| | | |
|---|---|---|
| *d1* | Rapamycin (Sirolimus) | a specific mTOR inhibitor |
| *d2* | Cladribine | adenosine deaminase inhibitor for U266, RPMI8226, and MM1.S cells |
| *d3* | Cetirizine DiHCl | an antihistamine. |
| *d4* | Venlafaxine HCl | Vearylalkanolamine serotonin-norepinephrine reuptake inhibitor (SNRI) |
| *d5* | Apixaban | selective, reversible inhibitor of Factor Xa |
| *d6* | Methylprednisolone | synthetic glucocorticoid receptor agonist |
| *d7* | Betaxolol | selective beta1 adrenergic receptor blocker |
| *d8* | Tolterodine tartrate | tartrate salt of tolterodine that is a competitive muscarinic receptor antagonist. |
| *d9* | CP-91149 | selective glycogen phosphorylase (GP) inhibitor |
| *d10* | Azithromycin Dihydrate | acid stable orally administered macrolide antimicrobial drug |
| *e1* | Temsirolimus (CCI-779, NSC 683864) | specific mTOR inhibitor |
| *e2* | Doxorubicin (Adriamycin) HCl | antibiotic agent that inhibits DNA topoisomerase II and induces DNA damage and apoptosis in tumor cells. |
| *e3* | Cilostazol | cyclic nucleotide phosphodiesterase type 3 (PDE3) inhibitor |
| *e4* | Voriconazole | triazole derivative similar to fluconazole and itraconazole that acts by inhibiting fungal cytochrome P-450-dependent, 14-alpha-sterol demethylase-mediated synthesis of ergosterol. |
| *e5* | Cefoperazone | cephalosporin antibiotic for inhibition of rMrp2-mediated [3H]E217βG |

| | | |
|---|---|---|
| *e6* | Ramipril | angiotensin-converting enzyme (ACE) inhibitor |
| *e7* | Ibutilide Fumarate | a Class III antiarrhythmic agent that is indicated for acute cardioconversion of atrial fibrillation and atrial flutter of a recent onset to sinus rhythm by induction of slow inward sodium current, which prolongs action potential and refractory period of myocardial cells. |
| *e8* | Quinapril HCl | an angiotensin-converting enzyme inhibitor |
| *e9* | PH-797804 | pyridinone inhibitor of p38α |
| *e10* | Tofacitinib (CP-690550) Citrate | inhibitor of JAK |
| *f1* | Crizotinib (PF-02341066) | inhibitor of c-Met and ALK |
| *f2* | PFI-1 (PF-6405761) | selective BET (bromodomain-containing protein) inhibitor for BRD4 |
| *f3* | Doxazosin Mesylate | selectively antagonizes postsynaptic α1-adrenergic receptors |
| *f4* | Ziprasidone HCl | dopamine and serotonin (5-HT) receptor antagonist |
| *f5* | Ketoprofen | dual COX1/2 inhibitor |
| *f6* | Nifedipine | dihydropyridine calcium channel blocker |
| *f7* | PF-4708671 | cell-permeable inhibitor of p70 ribosomal S6 kinase (S6K1 isoform)selectivity for S6K1 than MSK1 and RSK1/2, respectively. First S6K1-specific inhibitor to be reported. |
| *f8* | Miglitol | oral anti-diabetic drug. |
| *f9* | Dacomitinib (PF299804, PF299) | a potent, irreversible pan-ErbB inhibitor, mostly to EGFR |

| | | |
|---|---|---|
| *a1* | Gabapentin HCl | Gabapentin HCl is a GABA analogue, used to treat seizures and neuropathic pain. |
| *b1* | Varenicline Tartrate | Varenicline Tartrate is a nicotinic AChR partial agonist |
| *g1* | SU11274 | selective Met inhibitor |
| *g2* | Letrozole | third generation inhibitor of aromataseluteinizing hormone (LH), follicle-stimulating hormone (FSH), or androstenedione and does not affect normal urine electrolyte excretion or thyroid function in clinical studies. |
| *g3* | Etodolac | nonsteroidal anti-inflammatory drug (NSAID) and a COX inhibitor |
| *g4* | Orantinib (TSU-68, SU6668) | anti-PDGFR autophosphorylation ,also strongly inhibits Flk-1 and FGFR1 trans-phosphorylation |
| *g5* | Dofetilide | selective potassium channel ((hERG)) blocker |
| *g6* | Amlodipine | Amlodipine is a long-acting calcium channel blocker |
| *g7* | Bazedoxifene Acetate | selective estrogen receptor modulator (SERM). |
| *g8* | Clindamycin palmitate HCl | a water soluble hydrochloride salt of the ester of clindamycin and palmitic acid and a lincosamide antibiotic. |
| *c1* | Palbociclib (PD0332991) Isethionate | selective inhibitor of CDK4/6 |
| *d1* | Clindamycin Phosphate | lincosamide antibiotic for Plasmodium falciparum |

| | | |
|---|---|---|
| *h1* | PF-04217903 | selective ATP-competitive c-Met inhibitor |
| *h2* | Celecoxib | selective COX-2 inhibitor |
| *h3* | Fluconazole | fungal lanosterol 14 alpha-demethylase inhibitor |
| *h4* | Alprostadil | a Prostaglandin Analog and Prostaglandin E1 Agonist. |
| *h5* | Glipizide | Glipizide is used to treat high blood sugar levels caused by a type of diabetes mellitus called type 2 diabetes. |
| *h6* | Sulbactam | Sulbactam is a beta-lactamase inhibitor |
| *h7* | PD128907 HCl | selective dopamine D3 receptor agonist |
| *h8* | PF-04929113 (SNX-5422) | selective HSP90 inhibitor |
| *h9* | PF-5274857 | selective Smoothened (Smo) antagonist |

**Supplementary table 2. Sequences of siRNAs used in this paper.**

| Name | Sense | Antisense |
|---|---|---|
| si-CAPN1#1 | CAUGGAUCGUGAUGGCAAUTT | AUUGCCAUCACGAUCCAUGTT |
| si-CAPN1#2 | GGAGUUGUGACCUUUGACUTT | AGUCAAAGGUCACAACUCCTT |
| si-Beclin-1#1 | CCCAGGAGGAAGAGACUAATT | UUAGUCUCUUCCUCCUGGGTT |
| si-Beclin-1#2 | GGUCUAAGACGUCCAACAATT | UUGUUGGACGUCUUAGACCTT |
| si-Beclin-1#3 | GGACAACAAGUUUGACCAUTT | AUGGUCAAACUUGUUGUCCTT |
| si-ATG3-1#1 | GCGGAUGGGUAGAUACAUATT | UAUGUAUCUACCCAUCCGCTT |

| | | |
|---|---|---|
| si-ATG3-1#2 | GAGGCUACCCUAGAUACAATT | UUGUAUCUAGGGUAGCCUCTT |
| si-ATG3-1#3 | GCUUCAAGAUUGCUCAGCATT | UGCUGAGCAAUCUUGAAGCTT |
| si-ATG5-1#1 | GGACGAAUUCCAACUUGUUTT | AACAAGUUGGAAUUCGUCCTT |
| si-ATG5-1#2 | CCAUCAAUCGGAAACUCAUTT | AUGAGUUUCCGAUUGAUGGTT |
| si-ATG7-1#1 | CCAACACACUCGAGUCUUUTT | AAAGACUCGAGUGUGUUGGTT |
| si-ATG7-1#2 | CCAACAUCCCUGGUUACAATT | UUGUAACCAGGGAUGUUGGTT |
| si-control | UUCUCCGAACGUGUCACGUTT | ACGUGACACGUUCGGAGAATT |

**Supplementary Table 3. DNA primer sequences for quantitative real-time PCR.**

| **Name** | **Coding** | **Anticoding** |
|---|---|---|
| PD-L1 | TGGCATTTGCTGAACGCATTT | TGCAGCCAGGTCTAATTGTTTT |
| GAPDH | GGAGCGAGATCCCTCCAAAAT | GGCTGTTGTCATACTTCTCATGG |
